# Rapid spatial cognition in mice, with and without neocortex and hippocampus

**DOI:** 10.64898/2026.08.30.747945

**Authors:** Jieyu Zheng, Rogério Guimarães, Zeynep Turan, Anwesha Das, Jennifer Y. Hu, Katelyn Sadorf, Pietro Perona, Markus Meister

**Affiliations:** Division of Biology and Biological Engineering, California Institute of Technology; Division of Engineering and Applied Science, California Institute of Technology

## Abstract

Rapid learning, memory, and generalization are often attributed to the neocortex and hippocampus. We examined these cognitive abilities in mice navigating a reconfigurable 3D labyrinth. Naïve wildtype mice solved a complex route within a few rewards, generalized to similar problems, and learned to favor a decision far removed from reward experience, all within a few hours. Computational modeling suggests that much of the learning occurs even before the first occurrence of reward, as the animal builds a model of the environment. Mutant mice lacking neocortex and hippocampus retain all these cognitive abilities, and remember the task over multiple weeks. Their main deficit is a delay of the learning curve from inefficient exploration. In a few hours of exposure to the task, the subcortical brain has little opportunity to reorganize in response to the challenge. We conclude that rapid spatial learning in mice does not strictly require circuits of the hippocampus or neocortex.

## 1 Introduction

Biological intelligence turns limited experience into useful behavior across time and context. A single situation can impose many demands at once: the animal may need to infer what mattered from only a few informative events, refine that knowledge as experience accumulates, identify which decisions in an extended action sequence were responsible, preserve what it learns over time, and reuse it in new but related situations. These demands of rapid cognition are conventionally partitioned into separate faculties — learning (rapid extraction, refinement, and credit assignment), memory (retention), and generalization (transfer to new settings) — studied behaviorally in animals [1–4] and formalized in machine learning as few-shot learning, transfer learning, and meta-learning [5, 6].

Rodent studies provide strong evidence for many capabilities of rapid cognition. One-trial fear conditioning and avoidance learning show that a single experience can support lasting memory, sometimes over long intervals [7, 8]. In spatial tasks such as the Morris water maze, rodents learn a new goal location after one informative episode and use that memory immediately on the next trial [9, 10]. Prior experience in the water maze further accelerates learning: rats can acquire a new place within one to two trials each day [11]. More naturalistic maze studies show that mice can discover rewards and produce long, correct action sequences after only a few experiences [12, 13].

These findings have shaped strong expectations about the underlying neural systems. The hippocampus has been a central candidate for spatial learning and long-term memory [14, 15]. Neocortex, the evolutionarily most recent six-layered cortex, has been linked to the extraction of structured knowledge, with prefrontal cortex in particular implicated in abstraction, rule learning, and cognitive flexibility [16, 17]. While this literature does not establish that hippocampus and neocortex are required for every form of rapid cognition, it has made them the default framework through which such behavior is interpreted [18].

A major limitation of the existing literature is that the behavioral components of rapid cognition have been studied in isolation, each in its own dedicated task. Yet in natural behavior these processes are interleaved rather than separable, so how they work together cannot be inferred from any one of them alone. The central question is therefore not whether rodents can exhibit these abilities in principle, but how they are related. For example, latent learning means that experience can be acquired before any reward [19, 20], but how such pre-reward learning supports subsequent few-shot learning is poorly characterized. Credit assignment is well studied, yet how credit reaches a decisive choice point that is remote from reward in both space and time remains unclear [21, 22]. Rodents can form generalizable structure such as schemas under dense reinforcement [23, 24], but what rule, if any, an animal can extract from exploration before reward is ever encountered, and then carry into a new task remains unknown. Last but not least, it is an open question whether hippocampus and neocortex are strictly required for these abilities, rather than merely the neural substrates in which they are usually observed and recorded.

Here we bring these components of rapid cognition together using the Manhattan Maze, a three-dimensional navigational task with an explicit graph structure. Its branching, graph-like layout echoes natural rodent environments such as the underground burrows mice inhabit [25, 26]. This promotes species-typical exploration while keeping the task structure fully defined. Mice explore the maze freely and improve with experience, producing behavior that unfolds as a sequence of discrete turns and choices. Within this single task, latent learning, credit assignment, and generalization can be observed together in the same animals, over a few sessions lasting mere hours.

The first set of experiments measures the performance of wildtype mice across several dimensions of rapid cognition: how quickly mice extract efficient action sequences from sparse rewards; whether subsequent improvement is uniform or unfolds in distinct phases over the first hours of experience; how fast credit is assigned when efficient navigation depends on a decision remote from reward; how much learning is retained across days; and how much prior experience transfers to new maze configurations.

The second part of the study tests the brain systems long placed at the center of these behaviors. Mice lacking both neocortex and hippocampus reveal whether, and to what extent, spatial navigation depends on these structures. Situated in the Manhattan Maze, this test measures how much behavior requires the canonical hippocampal–neocortical framework, how much can arise from remaining subcortical structures, and which stage of learning the loss affects.

Finally, we contrast the behavioral findings with the predictions of candidate learning algorithms including model-free and model-based reinforcement learning (RL). The discrepancies that emerge underscore the study’s broader aim: to provide a reference point for rodent cognition and to assist the search for algorithms that learn from only a few experiences, a standing gap for NeuroAI [27].

## 2 Results

### 2.1 Graph-based task design of the Manhattan Maze

We designed an 11×11 two-layered *Manhattan Maze* (Fig. 1A and Fig. S1A-B) inspired by the grid layout of Manhattan Island (Section 5.4). Perpendicular “streets” and “avenues” were separated into two layers, connected via a changeable mask with holes that determine inter-layer access (Fig. S1C). This modular design allows 2^11^*^×^*^11^ ≈ 10^36^ unique configurations connecting the 22 corridors. In experiments, mask swaps alter the connectivity within seconds, while keeping the maze in the same physical space (Video V1). Thus, each mask change modifies the graph structure of the maze without requiring animals to learn an entirely new place representation.

**Figure 1:**
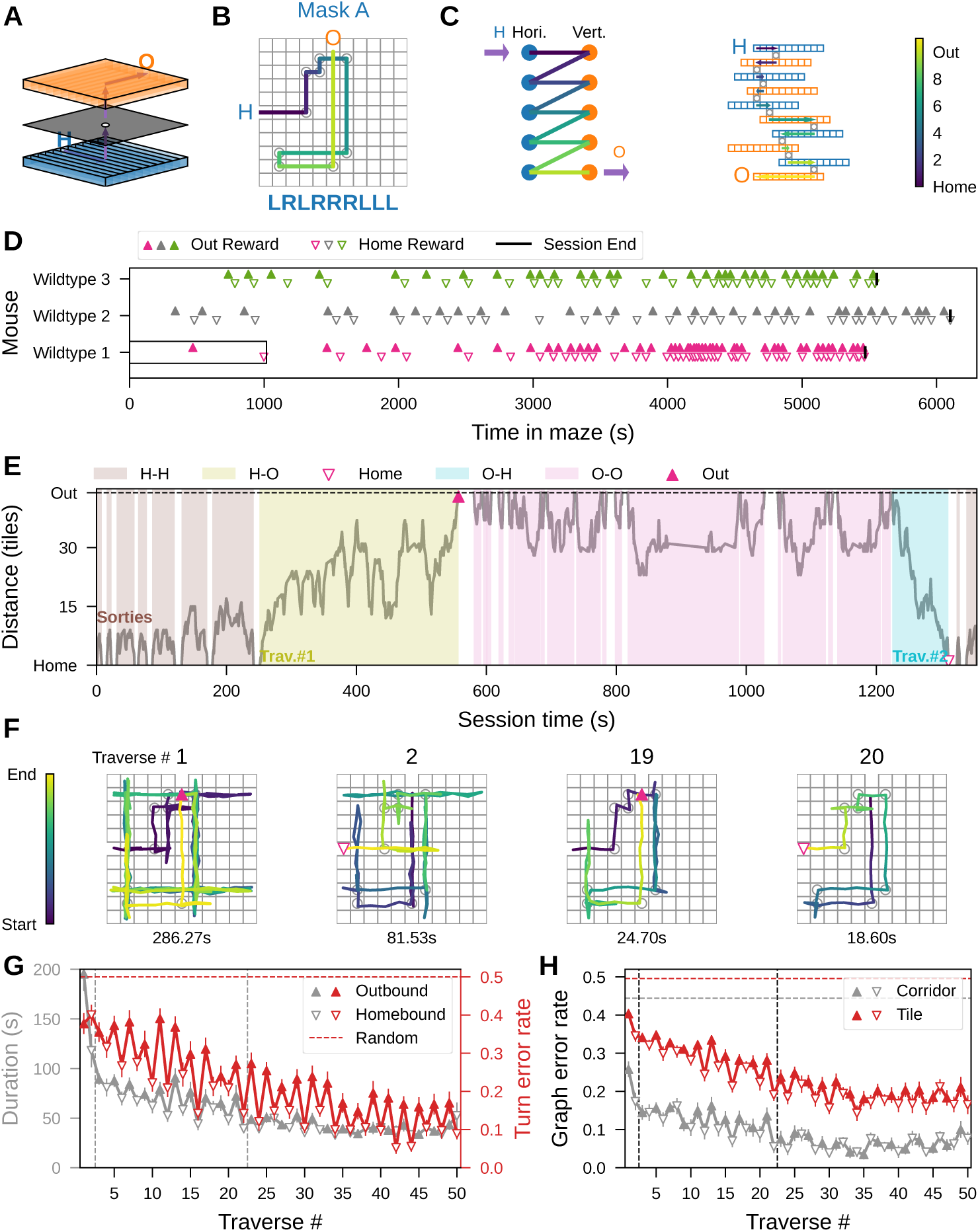
Rapid learning of the first map. **A-C**. The Manhattan Maze. **A**. The 2 × 11 × 11 Manhattan Maze: a reconfigurable maze pairing a bottom tray (blue) of 11 horizontal corridors with an inverted top tray (orange) of 11 vertical corridors, joined by a *mask*, a plastic sheet whose holes connect specific top and bottom corridors. Gray shows the simplest mask, Mask O, with one hole. Two water ports sit in the peripheral walls: “Home” (H, blue, bottom) and “Out” (O, orange, top). **B**. Top view of the maze with Mask A. Square tiles, corridor intersections where a hole may be placed and provide units of distance for our analysis. Mask A has 9 holes (circles) strategically placed, and the shortest path from Home to Out traverses 10 corridors is a 9-turn sequence (LRLRRRLLL, L=left/R=right) in egocentric coordinates. **C**. (Left) The corridor graph of the maze with Mask A (blue, horizontal corridors; orange, vertical). (Right) each corridor’s 11 tiles, segmented by holes into runs, turns and dead-ends. **D-H.** Learning the first mask. **D.** Rewards over time for three wildtype mice entering for the first time in the Mask A maze. Black vertical bars, session ends. Black rectangle, the time interval (sum of the colored blocks) plotted in E for mouse Wildtype 1. **E.** First exploratory bouts of Wildtype 1 (from D), shown as tile distance from Home over time. We indicate in color the time intervals blocks where the mouse proceeds in different directions: H-H (exploration around home), H-O (outbound), O-O (exploration around out port), O-H (homebound). Blank blocks, stasis at ports or home cage, excluded from in-maze time. **F.** We call “traverse” an entire path from H to O or vice-versa. Traverses #1, 2, 19, and 20 of Wildtype 1. Triangles mark traverse endpoints; random jitter separates overlapping trajectories. **G.** Left axis (gray): traverse duration vs. traverse number (25 mice); right axis (red): turn error rate. Vertical dashed lines divide the three learning phases. Horizontal dashed line, chance error rate (0.5). Error bars, mean ± standard error (SE) across animals. **H.** Corridor error rate (gray) and tile error rate (red) over traverse number (25 mice, mean ± SE). Dashed lines, memoryless-walker nulls (corridor 0.44, tile 0.50).

The maze has two access holes: one is connected to a mouse cage, which includes the *Home* water port; the other is connected to the *Out* water port. The water ports release a drop of water after a nose poke, but only in alternation, requiring the animal to run back and forth through the maze (Section 5.5.1). All mice started completely naïve to the maze and reward system (Section 5.5). After ∼20 h of water deprivation, the mice were introduced to the arena under complete darkness, while videos were recorded under infrared lights (Fig. S1A).

We describe the mouse trajectories by *bouts* that start and end at water ports (Section 5.6.2). Bouts between different ports (H-O and O-H) are termed *traverses*; bouts between the same ports (H-H and O-O) are *sorties*. A session is defined as a continuous period during which the mouse experienced a single, fixed mask configuration (Section 5.5). It begins with H-H (sorties), followed by the first H-O (*outbound traverse*) to a reward at the O port (Fig. 1E and Video V2). Then the animal performs O-O (sorties) and an O-H (*homebound traverse*). The session cycles through outbound (H-H → H-O) and homebound (O-O → O-H) *journeys*, each comprising all the sorties and the subsequent traverse from the same port. Because of the alternating reward schedule, only traverses were rewarded at the ending port (Section 5.5.1).

All mice were first acclimated to Mask O, a simple one-hole configuration used for learning the alternating reward delivery system (Fig. S1B). Within 2 h, mice learned to rapidly alternate between the ports and obtain water rewards (Fig. S1D). After this training, we introduced the mice to the learning challenges below.

### 2.2 Rapid learning of a 9-turn maze in three distinct phases

In the first set of experiments, the mice were required to learn three masks designed as *path graphs*: 10 corridors are connected by holes in a single chain between H and O (Section 5.4.1, Fig. 1B). The path graph can be solved by a purely local rule without a spatial map: never returning through the hole just crossed. However, each hole junction presents four choices (going forward and backward in the same corridor, or climbing through the hole for left or right turn), with only one turn leading to reward and the other three to dead ends or to the previous hole (Fig. S1C). Thus, the shortest path between the two water ports requires nine correct four-way decisions (Fig. 1C). This design is similar to Tolman’s T-alley maze [20] but increases complexity with four (rather than three) options at each intersection.

To follow the learning process, we employed several metrics (Section 5.6.3). At the coarsest level, *reward interval* measures the time between successive rewards, including the full journey between water ports. The *traverse duration* measures travel time for goal-directed navigation between the ports. The *turn error rate* captures the accuracy of decisions at the holes. The *corridor error* counts mistakes in which the animal crosses into a corridor farther from the goal. Finally, the *tile error* counts steps away from the goal on a finer scale: a “tile” is a square the width of a corridor (Fig. 1C). This error includes wrong turns, overshooting past a hole, and backtracking. Both corridor errors and tile errors were normalized by the total steps in a bout, expressed as *graph error rates* with a chance level near 0.5 (Section 5.7.1).

In the Mask A maze on Day 1, 25 mice showed rapid improvement within the first 20 rewards. Reward intervals quickly decreased to ∼0.11 of the starting value (Fig. 1D and Fig. S1E). Early in the session, mice often required several minutes to complete a single pair of traverses (Fig. 1E,H), frequently entering dead ends and repeatedly scanning the same corridors (Fig. 1F). As learning continued, their trajectories became increasingly direct, and Traverse 22 required only ∼0.2 of the initial duration (Fig. 1F,G). This dramatic acceleration of traverses was only partly due to faster locomotion, as the running speed increased only ∼1.5× (Fig. S1F-H).

Looking more closely, traverse performance appeared to improve in three qualitatively distinct phases (Fig. 1G). In the first phase, duration dropped sharply: the second traverse was already ∼0.6 of the first, even though it followed the opposite route through the maze. In the second phase, lasting ∼20 traverses, both duration and turn error rate declined gradually. In the third phase, both these measures stabilized. Notably, homebound traverses were consistently less erroneous than the preceding outbound traverses, an effect that we traced to subtle asymmetries of the maze (Section 6.6).

The graph error rates for corridor and tile choices followed a similar time course (Fig. 1H). Interestingly, both graph error rates started out considerably below chance (Fig. 1H), indicating that the animals had already learned to navigate the graph more efficiently even before collecting the first reward (latent learning). Indeed, the average mouse acquired a substantial forward bias along the way to the very first reward, outperforming a memoryless random walker at a quarter of the journey (Section 5.7.1 and Fig. S1I). Neither of these error rates reached zero, indicating that the average mouse did not converge on the shortest path through the maze.

### 2.3 Overnight memory and generalization to new mazes

On Day 2, following an overnight rest after learning Mask A, the mice were introduced to two new configurations: Mask B and Mask C (Section 5.5.5, Fig. 2A). Mask B has the same egocentric turn sequence as Mask A but uses different corridors, while Mask C has an entirely different turn sequence (Section 5.4.1). The two new masks were designed to test whether a familiar turn sequence helps in learning a new environment. The 25 mice were divided into six groups, each tested with a specific mask order following a permuted “XYXZ” design (Section 5.5.5, Fig. 2B). The four sessions were numbered sequentially as Day 2.1 to Day 2.4.

**Figure 2:**
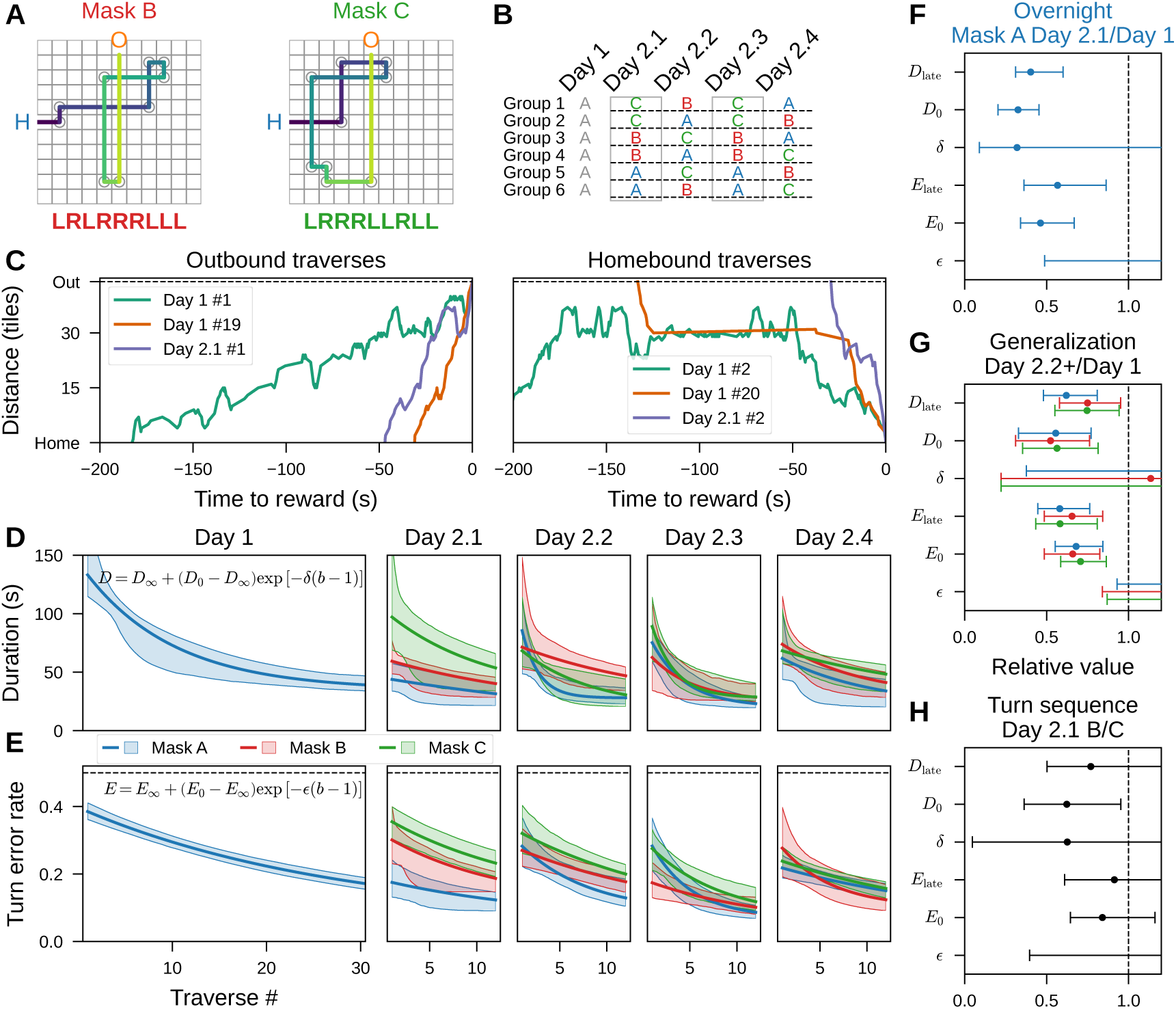
Overnight memory and generalization on Day 2. **A.** Mask B and Mask C. Mask B preserves Mask A’s turn sequence (LRLRRRLLL); Mask C differs at four turns (LRRRLLRLL). **B.** Experimental schedule: six groups of mice were assigned different sequences of mazes: one on Day 1 and four on Day 2. Each sequence was a different permutation of Masks A, B, and C. In all groups the masks on Day 2.1 and Day 2.3 were identical. **C.** Example outbound (left) and homebound (right) trajectories of one mouse from Group 5 in B. Left: the first outbound traverse on Day 1 (teal; Day 1#1), the one after 20 rewards on Day 1 (orange; Day 1#19), and the first on Day 2.1 (purple; Day 2.1#1). Right: the corresponding homebound traverses (Day 1#2, Day 1#20, Day 2.1#2), in the same colors. Only the last 200 seconds of the traverses are shown. **D.** Exponential fits to traverse duration across the two-day experiment, by mask and session. Lines and bands, median and 95% confidence intervals. **E.** Exponential fits to turn error rate, in the same format. Dashed line, chance error rate (0.5). **F-H.** Relative exponential-fit parameters (median ratio with 95% confidence interval; see text): initial value (*D*_0_, *E*_0_), learning rate (*δ*, *ɛ*), and late in-range performance (*D*_late_, *E*_late_). **F.** Day 2.1 Mask A relative to Day 1 Mask A (8 mice, overnight retention). **G.** Day 2.2, Day 2.3, and Day 2.4 relative to Day 1, grouped by mask (generalization effect). **H.** Day 2.1 Mask B relative to Day 2.1 Mask C (turn-sequence effect).

To quantify the learning curves across these conditions, we applied exponential curve fits to the traverse durations (*D*) and turn error rates (*E*) for each session as a function of traverse number (*b*, integer starting from 1; Section 5.6.4 and Fig. S3A-B). Each curve is described by three parameters: initial performance (*D*_0_*, E*_0_); asymptotic performance (*D_∞_, E_∞_*) or late performance (*D*_late_, *E*_late_ for weak asymptote identifiability; Section 5.6.5); and the exponential decay rate (*δ, ɛ*). Parameter ratios across conditions estimate the effect sizes of overnight memory and generalization (Section 5.6.5).

In session Day 2.1, the mice retested on Mask A showed strong overnight retention. Their initial traverse duration was ∼0.3 of that on Day 1 (Fig. 2C-F, *D*_0_; Video V3), an improvement also evident in shorter reward intervals (Fig. S2A). The initial turn error rate was ∼0.5 of that on Day 1 (Fig. 2F, *E*_0_; Fig. S2E and Fig. S3C). Mice performed better on Day 2 than on Day 1 well beyond the initial traverses, with *D*_late_ reduced to ∼0.4 and *E*_late_ to ∼0.6 relative to Day 1. This overnight advantage was specific to the previously trained mask. Within Day 2.1, mice on the repeated Mask A outperformed those on newly introduced Masks B or C on turn error rate (Fig. 2E; Fig. S3).

The subsequent sessions Day 2.2 to Day 2.4 tested the ability to generalize maze learning to new masks. Across these sessions, the mice showed similar learning curves, whether or not they had seen that specific mask before (Fig. 2D,E). Both duration and turning error rate started considerably lower than in the naïve session on Day 1 (*D*_0_ to ∼0.6 and *E*_0_ to ∼0.7 of the Day 1 values; Fig. 2G). Both performance measures continued to improve during the session (*D*_late_ to ∼0.7 and *E*_late_ to ∼0.6 of the Day 1 values; Fig. 2G and Fig. S3). In fact the exponential learning rates (*δ* and *ɛ*) were similar to the values on Day 1. Notably, the corridor error rate on the first traverse in a new maze was only ∼0.6 of the Day 1 value and changed little within sessions (Fig. S2G). These changes show that mice bring experience with one maze to bear on solving a different one, yielding improvements in performance on the order of 1.3× to 1.9×.

What specific information did the mice carry from Day 1 to Day 2? The comparison between Masks B and C tests whether mice reused the egocentric turn sequence learned in the Mask A maze. On Day 2.1, mice performed better in the Mask B maze — which recapitulates Mask A’s turn sequence — than in the Mask C maze, taking less time and making fewer errors on the traverse (*D*_0_ reduced to ∼0.6; Fig. 2E, H; Fig. S2E). Over the following sessions (Day 2.2 to Day 2.4), this bias in favor of the turn sequence of Mask A disappeared (Fig. 2G, Fig. S2), as the performance on Mask C improved. In short, mice briefly benefited from a specific turn sequence encountered on Day 1, but the accumulating experience with subsequent mazes rapidly outweighed that benefit.

### 2.4 Rapid credit assignment to a remote decision point

Next, Mask D was introduced as a qualitatively distinct task from the 9-turn masks, designed to test how animals assign credit to specific decision points in a graph. Mask D consists of two densely connected corridor subgraphs, or bicliques, joined by bottlenecks (Fig. 3A; Section 5.4.1). The bicliques create many locally valid routes, whereas the bottlenecks are high-value decision points because they provide the only routes between bicliques. The central bottleneck poses a remote credit-assignment problem: it is sandwiched between the two bicliques and distant from reward delivery, so successful reward discovery provides only delayed feedback about the importance of this location. Seven mice, trained only in the Mask O maze and naïve to all other masks, were exposed to Mask D. The analysis below focuses on the six mice that obtained rewards in that maze.

**Figure 3:**
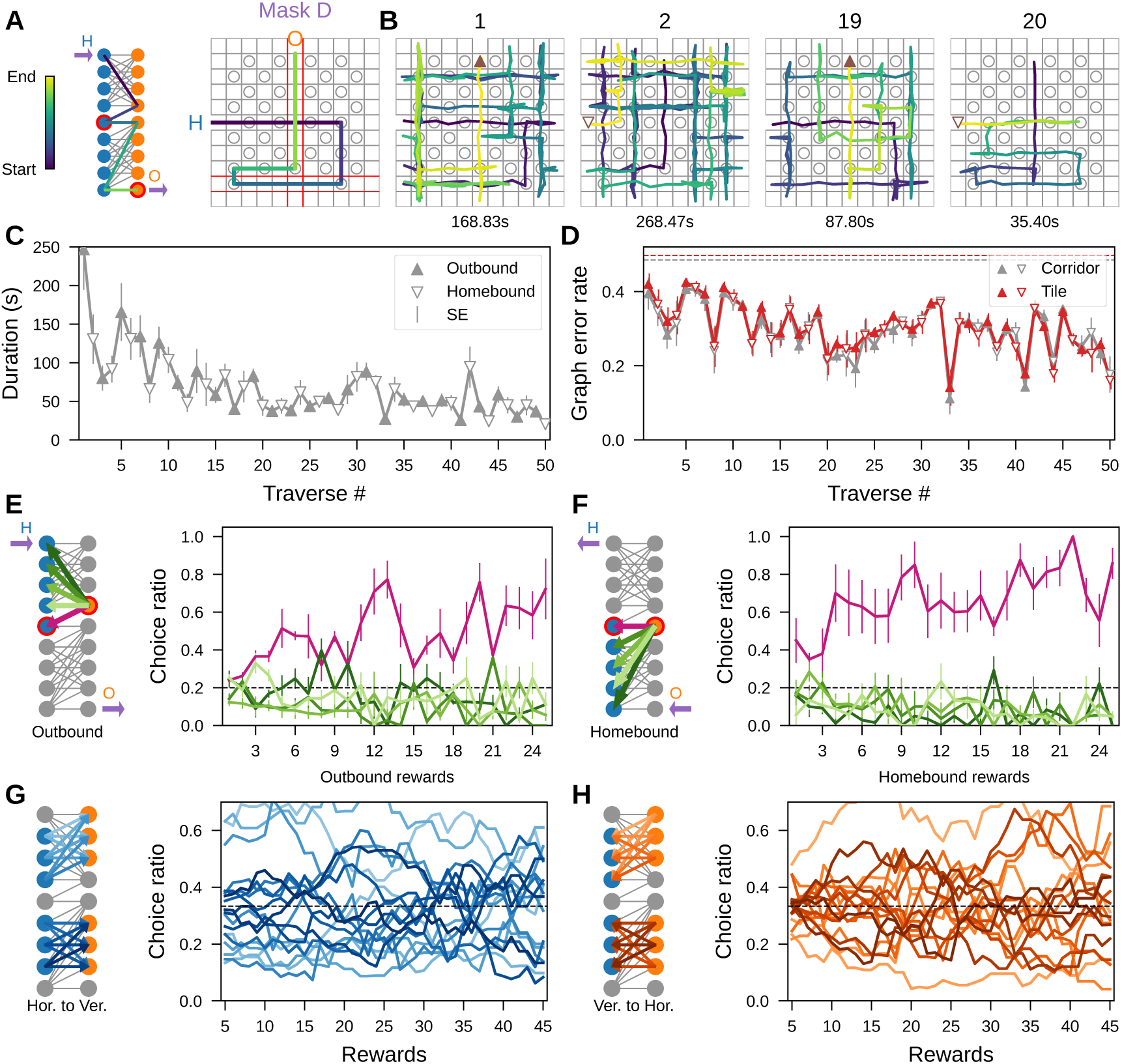
Rapid credit assignment to the bottleneck structure. **A.** (Left) Corridor graph of Mask D. (Right) top view. The shortest outbound path passes through two bicliques and bottlenecks (red). **B.** Traverses # 1, 2, 19, 20 of a mouse. **C.** Traverse duration vs. traverse number (6 mice, mean ± SE across animals). **D.** Corridor (gray) and tile (red) error rates over traverse number (mean ± SE). Dashed lines, chance error rates (corridor 0.49, tile 0.50). **E-H.** Preference towards the bottleneck. **E.** (Left) in the outbound direction, taking the bottleneck (pink arrow to the red circle) is the only exit from the first biclique at the gateway (orange), versus the four other corridors (green arrows). (Right) choice ratio for the transitions on outbound journeys. Dashed line, chance level (0.2). Error bars, mean ± SE. **F.** Same for the homebound direction: bottleneck (pink) vs. control (green) choice ratios at gateway (orange node). **G.** (Left) off-path transitions (blue arrows) from horizontal (blue circles) to vertical corridors (orange circles). (Right) Corresponding choice ratios for these transitions. Each line is population mean smoothed with a 10-reward moving average. Dashed line, chance level (1*/*3). **H.** Corresponding off-path vertical-to-horizontal transitions and choice ratios.

Learning in the Mask D maze followed the same trend as in the Mask A maze, showing a ∼5× improvement over the first 20 traverses. Traverse duration improved dramatically over the first two rewards, then more gradually, before stabilizing (Fig. 3B,C, Fig. S4A-D,E). Yet the animals never settled onto a fixed route, nor did they retrace their previous traverses (Fig. S5A-C). The similarity between traverse routes stayed low throughout the session (modified Jaccard similarity ∼0.2, Section 5.6.7 and Fig. S5B). The corridor and tile error rates tracked each other closely and declined to only ∼0.7 of the initial values (Fig. 3D), far too little to account for traverse duration (Fig. 3C). The persistently high graph error rates showed that animals were often off the optimal path (Fig. 3D). Late in learning, the animals found some very short routes, but they were never identical tile by tile (Fig. S5D, Video V4).

What did mice learn that let them accelerate their traverses? We found that the rapid learning was rooted in decisions near the bottleneck corridors (Fig. 3E,F; Section 5.6.6). When a mouse is at the gateway choice point adjacent to the bottleneck, how often does it pick the bottleneck as opposed to the 4 alternative options? That choice ratio for the bottleneck corridor began low but rose sharply within ∼3 rewards in the outbound direction (Fig. 3E) and remained above chance for the rest of the session. Notably, the first homebound journey already showed above-chance bottleneck choice, a hallmark of latent learning (Fig. 3F). In contrast, choice ratios for the other corridors stayed similar to one another and below chance. After crossing the bottleneck, mice were also less likely to reverse back into the previous biclique (Fig. S5E,F). For transitions among biclique corridors, no such pattern of preferences emerged (Fig. 3G,H). Together, these results indicate that mice were able to assign credit to the bottleneck link in the graph after just 1–3 rewards, while decisions within the bicliques remained flexible. Notably, the mice solved this assignment problem separately in both the outbound and homebound directions, even though these traverses were interleaved.

### 2.5 Acortical mice rapidly learned the maze after delayed reward discovery

In an attempt to connect these remarkable phenomena of rapid learning to brain function, we began with a gross perturbation of neuroanatomy: a mutant mouse that fails to develop its neocortex and hippocampus during embryonic development (Figs. S6 and S7). This model [28] exploits Emx1, a transcription factor selectively expressed in progenitors of projection neurons in the dorsal forebrain. Cre recombinase under control of the Emx1 promoter deletes a conditional allele of *Pals1*, a gene required for cortical development (Section 5.3.1). The lineage derived from these precursors, including projection neurons of the neocortex and hippocampus, is therefore absent. The mutants are homozygous conditional knockouts of *Pals1* (genotype *Pals1*^loxp/loxp^:*Emx1-Cre*^+^, “CKO” in Kim et al. [28]). We refer to these homozygous mutants as *acortical* mice, their Cre*^−^* siblings as *sibling* mice, and the C57BL/6J mice from previous sections as *wildtype* mice. Because both the Cre*^−^* siblings and the C57BL/6J mice are valid controls for the mutant line [29], and the two behaved indistinguishably where they can be compared directly (Fig. S8A), we pool them into a single *control* group for comparisons of learning Masks A, B, and C.

During training in the Mask O maze, acortical mice had difficulty obtaining rewards, largely due to repetitive scanning within corridors (Fig. S8A-C). Many were reluctant to enter the maze and needed human encouragement to crawl through the hole for the first time (Section 5.5); as a result, some mice failed to meet the learning criterion because they never explored the full maze or encountered the reward ports. Of all 18 acortical mice tested (Section 5.5.3), 100% learned Mask O with assistance (18 of 18), whereas 56% learned Mask A (9 of 16; Tables 1 and 2). To disentangle failed exploration from learning, the analyses below focus on the successful learners, defined as acortical mice that obtained at least 20 rewards in a single session. Four of seven acortical mice that met Mask A directly after Mask O learned it and are compared below with the pooled control group (27 mice).

**Table 1:** Learning results of all 18 acortical mice. A mask is considered learned only when the mouse obtained at least 20 rewards in one session. Masks not listed in either column were never presented to that mouse.

| Acortical mouse | Learned masks | Failed masks |
| --- | --- | --- |
| 830 | O | E, A, D |
| 780 | O, A, B, C, D |  |
| 777 | O, E | A, C, D |
| 735 | O, A, B, C, D |  |
| 683 | O, A, B, C, D |  |
| 655 | O, A, B, C, D |  |
| 589 | O | C, D |
| 585 | O, A, B |  |
| 078 | O, E | A, D |
| 077 | O, E |  |
| 076 | O, E, F, A, C | D |
| 073 | O, E, F, A, C, D |  |
| 072 | O, E | F, A, D |
| 070 | O, E, F, A | D |
| 069 | O, E, F | A, D |
| 067 | O, E, F, A | C, D |
| 049 | O, E | F, A, D |
| 047 | O | A, D |

**Table 2:**
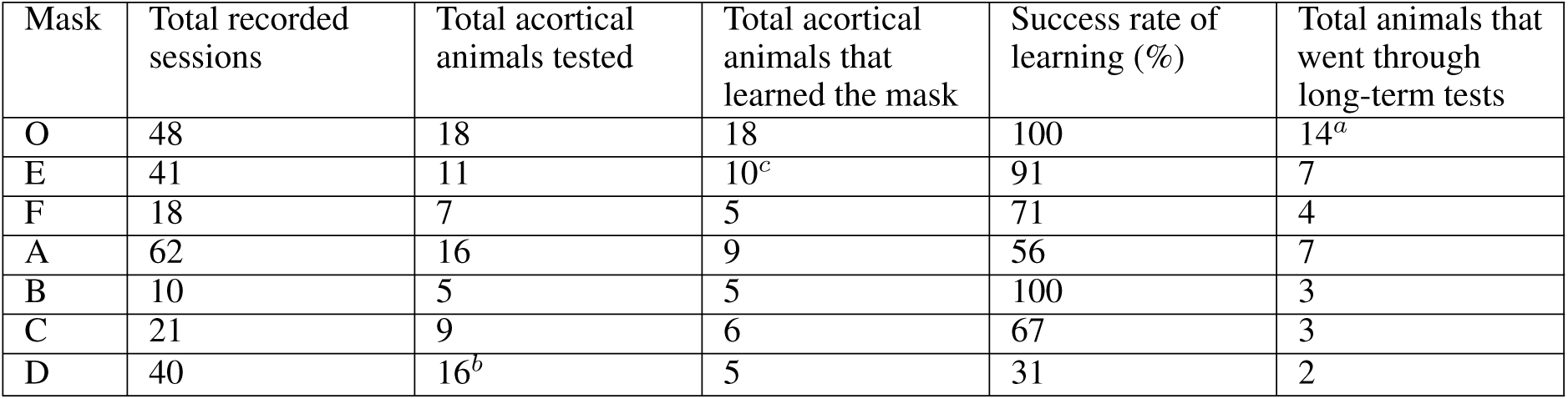
Number of successful mice for each mask. A mouse is considered to have learned a mask only after obtaining at least 20 rewards in one session. Only a subset of the successful animals were tested for long-term memory on the same masks. Success rate is computed over all animals tested on that mask.

The four successful acortical mice took ∼2.4× as long as control mice to obtain the first four rewards (Fig. 4A). This delay was not a motor impairment: the mice traversed the maze marginally faster than control mice (Fig. S8D). Instead, the delay reflected two stereotyped but inefficient behaviors. First, the mice performed excessive short sorties, repeatedly dashing back and forth between a port and the adjacent corridor (Fig. 4B, Video V5). Early in learning, acortical mice made ∼3.8× as many sorties as control mice (Fig. S8E). Second, the mice frequently scanned a corridor back-and-forth without passing through holes (Fig. 4B-D). Acortical mice traveled ∼1.5× as many tiles per corridor during the first two journeys (Fig. 4C). Even during subsequent learning, tile error rate stayed above that of control: reaching ∼1.2× the control value (Fig. S8F).

**Figure 4:**
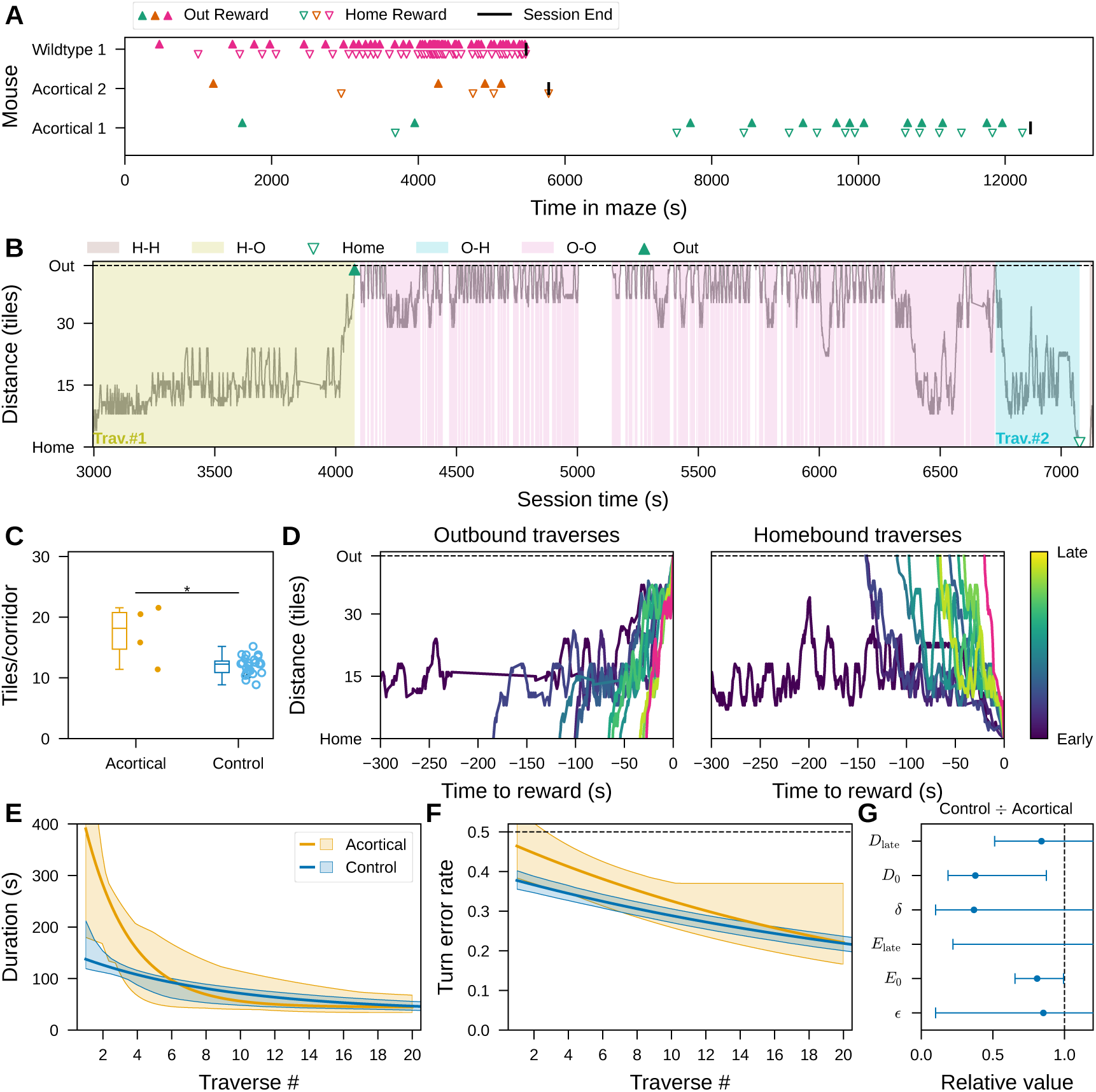
Acortical mice rapidly learned the maze after delayed reward discovery. **A.** Rewards over time for two acortical mice (teal, orange) and one wildtype mouse (pink, same as in Fig. 1D-F). **B.** The first two traverses (olive, outbound; cyan, homebound) and the O-O sorties between them (pink) of acortical mouse 1 (teal) from A, shown as tile distance from Home over time. **C.** Average tiles traveled per corridor over the first two journeys for acortical (4 mice, orange) and control (27 mice, blue; dark blue filled dots, sibling; light blue open dots, wildtype). The asterisk shows a significant two-sided Mann-Whitney U test (*U* = 92, *p* = 0.023). **D.** The first 10 outbound (left) and homebound (right) traverses of Acortical 1, plotted as tile distance from Home. Early to late traverses are plotted from dark to light. Pink lines, the ninth (left) and tenth (right) traverses by Wildtype 1 (pink) in A. **E.** Exponential fits of traverse duration for acortical (4 mice, orange) and control (blue) mice. Lines and bands, median and 95% confidence intervals. **F.** As in E, for turn error rate. Dashed line, chance error rate (0.5). **G.** Relative parameter values of control to acortical mice. Error bars, median and 95% confidence intervals.

Despite the initial difficulties with repetitive scanning (Fig. S8G), the acortical mice improved rapidly. Though their forward bias was much lower than that of controls, it increased pre-reward (Fig. S8I). Their traverse durations approached those of control mice within ∼10 rewards (Fig. S8G). Sorties became less frequent (Fig. S8E), yielding shorter reward intervals (Fig. 4A). Traverses became more direct (Fig. 4D) and committed fewer tile errors (Fig. S8F,H). Eventually, their trajectories were nearly as short as those of a wildtype mouse (Fig. 4D and Video V6).

Learning curves for traverse duration and turn error rate (fit with the methods of Sections 2.3 and 5.6.4) confirmed that the largest impairment of acortical mice was in the initial performance. The initial traverse duration (*D*_0_) was ∼2.6× that of controls (Fig. 4E,G; Fig. S9A), whereas the late duration *D*_late_ and learning rate *δ* were comparable across genotypes. Likewise, the initial turn error rate (*E*_0_) was slightly elevated (Fig. 4F,G; Fig. S9B), yet the late (*E*_late_) and learning rate (*ɛ*) were similar. Together, these results indicate that acortical mice are most impaired during the first phase of reward discovery, leading to a delay in the learning curves. However, successful learners retain the capacity for rapid improvement, eventually reaching performance comparable to controls.

### 2.6 Acortical mice retained maze solutions over days

We next investigated whether maze solutions acquired without cortex and hippocampus can be retained. To test this, masks were reintroduced after intervals ranging from overnight to several weeks (Section 5.5.3). No mouse encountered Mask A during the interval, and nearly all intervals longer than one week contained no maze experience of any kind (Section 5.5.3).

Acortical mice maintained performance across breaks when tested again on the same masks. Overnight retention was first evident in the Mask O maze (11 mice). Although initial learning was slow, all acortical mice obtained rewards almost immediately on the next day, with reward intervals ∼0.2 of those at the beginning of Day 1 (Fig. S10A).

In the Mask A maze, the example mouse shown in Fig. 5A began Day 2 with traverse durations and turn error rates close to the levels reached at the end of Day 1. Performance continued to improve over subsequent sessions, and perfect turn sequences emerged after a 6-day gap, on Day 8. Even after longer gaps of three weeks to one month, mice re-entered the maze with short traverse durations and low error rates. The animals also moved faster, reaching ∼1.5× the speed from Day 1 (Fig. S10B,C). Early traverses on later test days no longer showed repetitive scanning (Fig. S10D).

**Figure 5:**
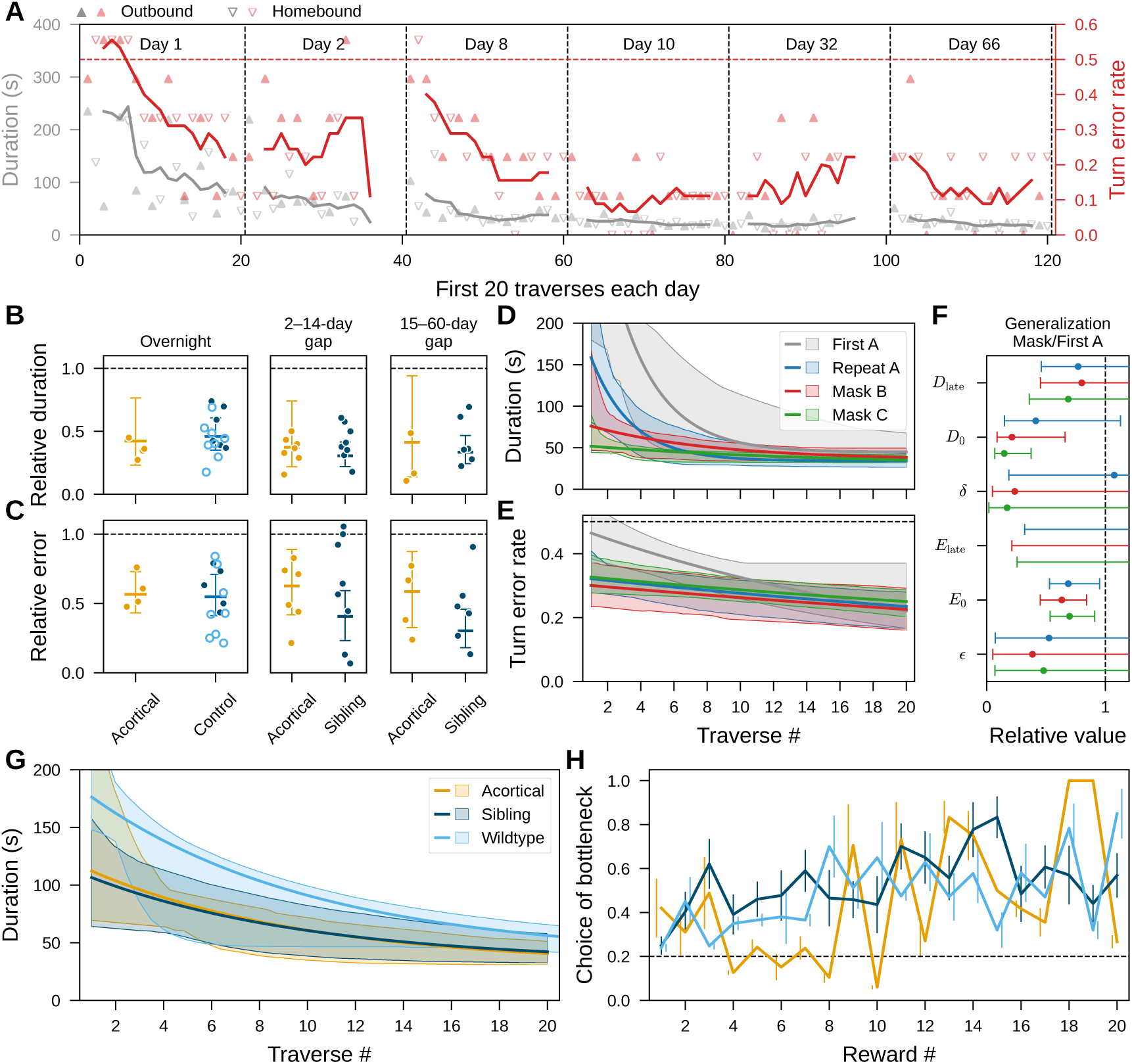
Acortical mice were capable of long-term memory and generalization. **A-C.** Long-term memory of Mask A. **A.** First 20 traverses of each session of an acortical mouse, showing traverse duration (left axis, gray) and turn error rate (right axis, red). Solid lines, moving averages over five traverses. Red dashed line, chance error rate (0.5). **B.** Relative mean traverse duration of acortical (6 mice, orange) and control mice. Columns, the gap since the previous Mask A session. The overnight column pools 8 sibling (dark blue) and 8 wildtype mice (light blue). Error bars, 95% confidence intervals of the session mean; dots, individual sessions. **C.** Relative mean turn error rate of the sessions in B. **D-F.** Generalization in 9-turn masks. **D.** Exponential fits of traverse duration of acortical mice for repeated Mask A (7 mice, blue), Mask B (5 mice, red), and Mask C (6 mice, green), relative to the initial Mask A (gray; identical to the orange acortical curve in Fig. 4E). **E.** Exponential fits of turn error rate (gray curve is identical to the orange acortical curve in Fig. 4F). Lines and bands, median and 95% confidence intervals. Dashed line, chance error rate (0.5). **F.** Relative parameter values for the repeated and new masks compared to the first Mask A. Error bars, 95% confidence intervals. **G-H.** Generalization in the Mask D maze. **G.** Exponential fits of traverse duration in the Mask D maze for acortical (5 mice, orange), sibling (8 mice, dark blue) and wildtype (6 mice, light blue). **H.** Choice ratio for the bottleneck corridors across all journeys (the pink transitions in the Fig. 3E-F schematics). The wildtype curve is equivalent to the two pink curves in Fig. 3E-F combined, interleaved by outbound and homebound. Dashed line, chance choice ratio (0.2)

To quantify retention, traverse duration and turn error rate were normalized to Day 1 performance (Section 5.6.3). Across genotypes, mice maintained shorter traverse durations (median ∼0.4) in later days than during the first encounter, even after intervals approaching 60 days (Fig. 5B). In acortical mice, turn error rates also remained consistently below Day 1 baselines, indicating retention of the learned turn sequence as well as faster traversal (Fig. 5C). At the population level, acortical traverse speed increased to ∼1.5× the Day 1 value during later days (Fig. S10C), indicating greater proficiency in navigation. Thus, once acortical mice acquired a maze solution, they retained it over long intervals, preserving short routes with a quality comparable to control mice.

### 2.7 Acortical mice generalized in new mazes

Because generalization depends on learning and retaining previously acquired solutions, the few-shot learning and long-term retention seen in acortical mice raise the possibility that they are also able to learn new maze configurations. This was tested by introducing additional masks after the animals had acquired an initial solution.

Generalization was first tested across the other 9-turn masks, Mask B and C (Fig. 2A), in mice that had previously learned Mask A or the intermediate Mask E (Sections 5.5.3 and 6.4). In these animals, learning a new mask was substantially faster than learning the first. Repetitive behaviors were reduced from the outset: shorter reward intervals (Fig. S11A) were caused by fewer sorties between rewards (Fig. S11B), and their first two traverse durations were only ∼0.2 of those during the first maze experience (Fig. S11C).

Generalization in acortical mice was expressed primarily as an improvement in initial performance. Both the traverse duration and the turn error rate started out much lower than in the initial maze (∼0.3 for *D*_0_, ∼0.7 for *E*_0_; Fig. 5D-F; Fig. S9C-E). Asymptotic performance and learning-rate parameters showed no consistent differences. Thus, as in wildtype mice, prior maze experience greatly improved the starting point of learning, rather than the subsequent learning rate (Fig. 2D,E; Fig. S3). Because Masks B and C were not presented in a fixed order, this experiment could not isolate transfer specific to the shared turn sequence (Fig. 5F).

Generalization was then tested in the Mask D maze, whose bottlenecks create a remote credit-assignment problem. Thirteen acortical mice and eight sibling mice that had previously learned at least one of Masks A, B, C, or E were introduced to Mask D. Five of the 13 acortical mice learned Mask D, whereas all eight sibling mice did. Among successful learners, both acortical and sibling mice outperformed wildtype mice encountering Mask D as their first complex mask (Fig. 5G): their first rewards were obtained ∼3× faster than by naïve wildtype mice (Fig. S12A), driven by fewer sorties and shorter traverses (Fig. S12B,C). As in the 9-turn masks, their improvement was mainly expressed in the initial-performance parameters (Fig. S9F,G). Within this small subset of successful learners, acortical mice showed a preference for the bottleneck corridors similar to wildtype and sibling mice (Fig. 5H), indicating an intact ability to assign credit to decisions remote from reward.

In summary, at least some acortical mice demonstrated the capacity for generalization, expressed as improved initial performance on new masks. Together with the few-shot learning and long-term memory results, this indicates a preserved suite of relevant cognitive functions in acortical mice, once they passed the exploration stage.

### 2.8 Candidate algorithms for few-shot spatial learning

How does the brain algorithmically support such rapid spatial learning in the Manhattan Maze, especially in complex configurations such as Mask D? A candidate algorithm should account for three observations: (1) few-shot learning within two rewards, matching the timescale of our measurements; (2) remote credit assignment to the bottleneck locations; and (3) plausible implementation by subcortical circuits, given that acortical mice also learned Mask D.

Two reinforcement-learning frameworks make contrasting predictions about how the bottleneck could be learned. The first is model-free reinforcement learning (RL), exemplified by the canonical TD(0) Q-learning algorithm (Section 5.7.3; 30). A Q-learning agent maintains a look-up table of values over state-action pairs. Each value is updated by a temporal-difference (TD) error, which is the discrepancy between the prediction of reward from the current state and the successor state. For a temporally local Q-learning agent, each update draws only on the immediately succeeding state, so reward information propagates backward by one step per rewarded episode. In the brain, the TD signal has been associated with the phasic activity of midbrain dopamine neurons [31].

The second framework is model-based RL, which builds an explicit map of the environment. It acquires the whole cognitive map without step-by-step reward-value propagation. We illustrate this with the Endotaxis model (Section 5.7.2 and 32), a neuromorphic implementation of model-based RL that respects biological learning rules. During exploration, the Endotaxis agent separately learns the cognitive map and the location of rewards on that map. The circuit includes point cells that represent the animal’s current location, map cells that learn the connectivity between locations, and goal cells that signal behaviorally relevant targets. As the agent moves, recurrent synapses between map cells are updated from the sequence of visited locations, so the map-cell network learns the local adjacency structure of the environment without reward experiences. When a reward is later encountered, the corresponding goal cell activates and propagates a goal signal through the learned adjacency structure. This produces an internal gradient that guides navigation by local comparisons among available paths, rather than an explicit global shortest-path calculation.

To compare the two models, agents implementing either Q-learning or Endotaxis learned by self-play on the Mask D corridor graph, starting from random walks. The sorties preceding the first outbound and homebound traverses carry no reward signal, so first-traverse performance for both models is identical and set by the memoryless random walker, with near 0.5 corridor error rate (Fig. 6A-B) and at chance (0.2) bottleneck choice ratio (Fig. 6C,D). However, once rewarded, the two models learn at markedly different rates. Q-learning’s updates propagate slowly, back through the second biclique, and its performance depends heavily on how the self-play traverses reach the reward. Walkers that pass directly through the biclique learn the bottleneck quickly, but those taking indirect routes take longer to propagate a value signal to the bottleneck. In comparison, Endotaxis learns the shortest path from a single rewarded experience, so both performance metrics become perfect at once.

**Figure 6:**
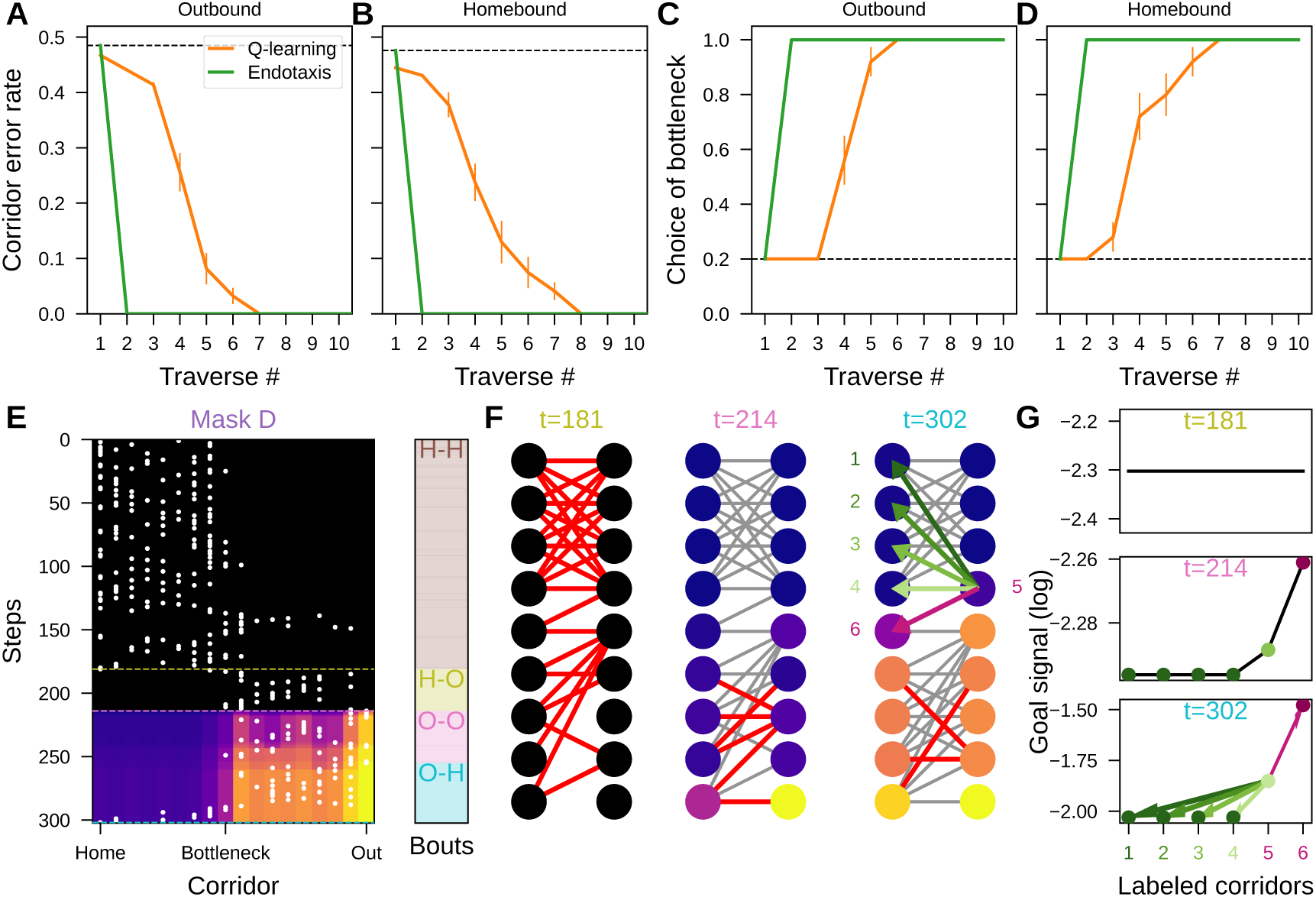
Endotaxis rather than Q-Learning acquires bottleneck in few-shot learning. **A-D.** Corridor error rate and bottleneck choice ratios by Endotaxis (green) and Q-learning (orange). **A.** Corridor error rate of outbound traverses. Dashed line, chance error rate. **B.** of homebound traverses. **C.** Choice ratios of the bottleneck in outbound traverses. Dashed line, chance choice ratio. **D.** in homebound traverses. **E-G.** Endotaxis model trained on the first two journeys of the mouse in Fig. 3B. **E.** Goal signal propagates through corridor visits, capped at 1 and log-scaled (lighter is higher). White dots, corridor visits. The black region of no signal ends when the first reward is obtained at the olive dashed line. Remaining dashed lines, the time points in F-G. Righthand strip, bout type along corridor steps. **F.** Goal signals (circles, lighter is higher) and learned adjacency on the Mask D corridor graph. Red edges, newly learned connections after the previous time point; gray edges, connections already learned at the previous time point. (Left to right) start of the first outbound traverse (*t* = 181 corridor steps), its end (*t* = 214), and the end of the homebound traverse (*t* = 302). Pink arrow, the bottleneck transition (corridor 4 to 5); green arrows, the control transitions, as in Fig. 3E. **G.** Goal signals at the bottleneck and control corridors for the three time points in F (*t* = 181, 214, 302). Labeled corridors and arrows follow F; dot colors match the corridors in F.

To examine this one-shot learning, we simulated Endotaxis with the first two journeys of the animal in Fig. 3B. Endotaxis learned the graph connections among corridors through the sorties, even before the first rewarded traverse (Fig. 6E-G). At this stage, the topological structure of the mask was already encoded by the map cells, but no goal signal had yet appeared in the corridors (Fig. 6E). Therefore Endotaxis trained on sorties alone cannot guide direct traverses to the goal. However, after the first reward, the goal signal appeared immediately and propagated through the pre-learned graph, creating a gradient through the bottleneck. Critically, the goal signal at the bottleneck was higher than at the control corridors within the bicliques (Fig. 6G). This pattern could immediately induce a directional preference toward the critical junction while treating the other corridors similarly (Fig. 3E-H).

In summary, both RL frameworks are biologically plausible, but only Endotaxis, not Q-learning, is able to capture few-shot learning. Endotaxis should be viewed as one possible explanation rather than a unique mechanism. It shows that the bottleneck structure can, in principle, be identified before the animal receives direct reward feedback about its importance. The proposed circuit does not depend on the trisynaptic recurrent loop of the hippocampus or the layered organization of the neocortex. It therefore offers a plausible algorithmic account of bottleneck learning, consistent with the preserved learning and generalization observed in acortical mice [32].

## 3 Discussion

In this study we quantified five aspects of spatial learning within a single task framework. In wildtype mice we observed: (1) few-shot learning of a novel route, reflected by the abruptly shortened traverses after the first one or two rewards (Fig. 1G, Fig. 3C); (2) gradual refinement of decisions, shown by corridor and tile errors declining over the next 20 rewards (Fig. 1H, Fig. 3D); (3) rapid credit assignment to a remote bottleneck, within 1–3 rewards (Fig. 3E,F); (4) retention of the same environment across days, including a successful turn sequence (Fig. 2C-F); and (5) generalization to novel configurations that does not rest on rote memory of those turns (Fig. 2G,H). Markers (1)–(3) trace how the map is assembled within a single session; markers (4) and (5) describe what becomes of it, in memory across days and in transfer to a maze the animal has never seen.

Removing the cortex and hippocampus dissociates these components. Acortical mice explored far less efficiently before their first reward (Fig. 4A-C). Once learning was underway, however, they converged to the traverse duration of controls (Fig. 4D-G), retained mazes across days (Fig. 5A-C) and generalized to new mazes as well as controls did (Fig. 5D-H). Fewer acortical mice reached criterion (Table 2), but those that did learned normally.

Notably, the one-shot improvement (1) and the structural component of generalization (5) can be traced to a single rule, while credit assignment to a remote decision point (3) requires a cognitive map. Together they constrain what class of learning algorithm can account for the behavior, explained in the next two sections. The final section discusses which of these functions are preserved in the absence of cortex and hippocampus.

### 3.1 Rapid rule learning and generalization

The initial phase of map learning included a dramatic one-shot effect (Fig. 1G,H): following the first outbound traverse, the second outbound traverse (Traverse #3) was ∼2× faster in duration. The first homebound traverse (Traverse #2) was already ∼1.7× faster as well, even though the animal had never run that route. Some knowledge must transfer from the outbound to the homebound trajectory, even though the two require entirely different decisions.

One source of this improvement was a reduction in the corridor error rate, with Traverses #2 and #3 falling to ∼0.7 and ∼0.6 of Traverse #1 (Fig. 1H). A corridor error occurs when the mouse reverses course, returning through the same hole it had just passed through. Instead, the mouse learned the rule “find a different hole from the one you just crawled through”. On all the path-graph mazes (A, B, C), this “forward rule” is beneficial: followed rigorously, it carries the animal through the maze in the minimal number of corridors (Table 3) and serves outbound and homebound traverses equally well. It also requires no knowledge of where the reward is and no association of specific places with turns, only a memory of the last hole. Finally, the forward rule is a prerequisite for the subsequent reward-driven learning of turns (Fig. 1G): without it, much of the animal’s experience would be gathered while traveling the wrong way, which would interfere with learning the proper association of locations with turns.

**Table 3:**
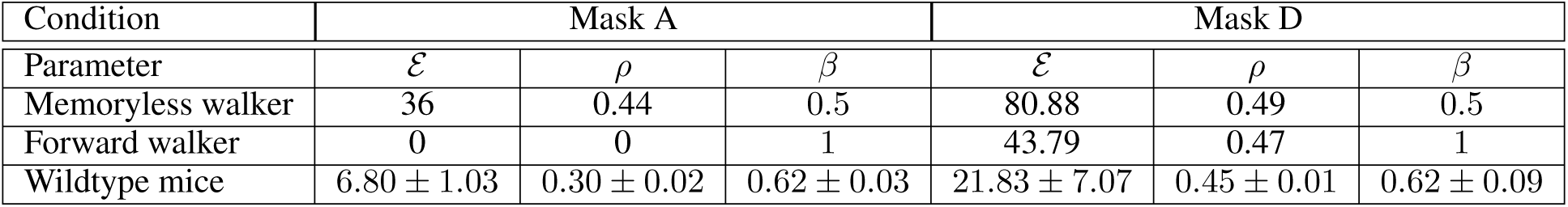
First-order Markov walker and mice on the first outbound traverse. For each mask the three quantities are compared across the memoryless walker (*β* = 0.5), the deterministic forward walker (*β* = 1), and the wildtype mouse population on its first home–out (H–O) traverse. *ε*, expected number of corridor errors; *ρ*, per-step corridor error rate; *β*, forward bias (model parameter for the walkers; empirical estimate 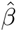 for the mice). Mouse entries 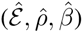 are mean ± SE across animals (Mask A, Fig. 1G, 25 mice; Mask D, Fig. 3C, 6 mice).

The animals acquired this bias over the course of the first journey (Fig. S1I): the empirical forward bias *β*^^^ rose from ∼0.35 during the unrewarded sorties and peaked at ∼0.6, a ∼1.7× increase. Crucially, it passed the memoryless walker’s value of *β* = 0.5 about a quarter of the way through the journey, well before any reward was encountered. The consequence for performance improvement is large. The memoryless walker scores a corridor error rate of 0.44 on a traverse (Section 5.7.1), whereas the average mouse was already at 0.26 on its very first traverse (Fig. 1H). No reinforcement-learning agent can assign value to this behavior before the first reward, whether it learns model-free or model-based (Fig. S13B,C,F). Moreover, the rule was acquired at every corridor in parallel as a global update, not one corridor at a time working backwards from the reward port (Fig. S13A,B).

The forward rule should not be confused with the “forward-going tendency” reported in the early literature on rats running mazes, where a preference for one route over another was attributed to the mechanical ease of executing different turns [33, 34]. Reversing at a T-junction requires a stop, a 180*^◦^* turn, and a re-acceleration, whereas running through the junction is easier. In the Manhattan Maze, by contrast, all the holes in a given corridor are mechanically equivalent; no feature favors the “forward” hole over the “reverse” hole, especially in early trajectories, where the animal often scans the entire corridor (Fig. 1F). The forward rule in the present study is therefore a purely cognitive choice, not a locomotor constraint.

Beyond the first maze, the forward rule also transfers knowledge to new ones. When the mice were confronted with the Mask B or C maze on Day 2, their corridor error rate started far below the memoryless level (Fig. S2G). Initial traverse duration improved ∼2× on both mazes (Fig. 2G), consistent with retention of the forward rule and its systematic application in the new maze. Compared to other phenomena of transfer learning, it is remarkable how fast this rule is acquired: within a single session of one or two hours, and largely before the first reward, rather than over many days of repeated sessions and thousands of trials [1, 24, 35]. The difference may lie in the special ecological significance of navigation rules, as opposed to the abstract challenges posed in other rule-learning tasks.

In sum, the forward rule is a clear example of structural knowledge obtained without reward, which then assists in reward-based learning.

### 3.2 Rapid learning of remote decisions

Mask D presented the animals with a special challenge: two heavily interconnected environments (bicliques) joined at a single bottleneck corridor. The “forward rule” above, with its limited memory, cannot solve this maze efficiently: a deterministic forward walker (*β* = 1) lowers the corridor error rate only from 0.49 to 0.47 (Section 5.7.1 and Table 3). It is easy to get lost within either biclique, so there is great value in finding the bottleneck, passing through it, and not reversing again. Remarkably, the animals came to prefer the bottleneck at the gateway after a single reward in the homebound direction and three rewards in the outbound direction (Fig. 3E,F). They solved the two problems alternatingly, on outbound and homebound traverses, even though these are separate gateway decision points with disjoint alternatives. Further, although the bottleneck sits only three or four corridors from the reward along the shortest path, the animal’s actual route between them was far longer: during early trajectories it took many wrong turns within the bicliques (Fig. 3B), such that tens of seconds and dozens of steps separated the bottleneck decision from its consequence. How can such remote decisions be learned so efficiently?

Under both broad solutions, model-free and model-based learning [30], the agent’s goal is to learn the value of the actions available in each state, for example at each hole of the maze. In model-free learning, the experience of reward raises the value of the immediately preceding action. Within biological neural circuits this can be implemented by strengthening the synapses that link the neural representations of states and actions, gated by a global signal that marks the reward: a three-factor synaptic update rule [31, 36]. Such updates can reach back in time to reinforce actions taken earlier, using synaptic eligibility traces [22]. Biological mechanisms of plasticity are thought to operate on time scales of about 1–2 s [21, 37]. Every experience of a reward, or of a state previously learned to have high value, can reinforce decisions reaching back no more than 2 s. In our experiments, the first reward was separated from the preceding bottleneck decision by tens of seconds to minutes (Fig. S4F), 10× to 100× beyond the 2 s window.

In model-based learning, by contrast, the agent has a map of the environment [20, 30]. On discovering a reward location, the agent can simply mark the associated state on the map and, on a future occasion, use the map to navigate a route to that state. This of course requires some means of learning the map in the first place. A specific form of the cognitive map suitable for neural implementation is the successor representation [38, 39], which can be learned as the agent explores, using biologically realistic synaptic update rules that do not depend on reward experience at all [32, 40]. By the time the agent finds the first reward, random exploration has already set up the cognitive map, and the reward location can be learned in one shot. We simulated a particular version of this scheme (Endotaxis [32]) using actual mouse trajectories (Fig. 6E-G). Before the first reward, the goal signal is uniformly at floor across the maze. Immediately afterwards, the value function has propagated through the entire map, with a monotone gradient rising from the control corridors through the gateway to the bottleneck, steep enough to guide the correct choice on the very next attempt (Fig. 6G).

Across both sets of mazes, our findings conflict with the trial-and-error framework that underlies model-free RL. Learning from reward as a teaching signal has long been a prominent account; but our data, together with previous reports of few-shot learning and sudden insight [13], are inconsistent with its key predictions. Trial and error predicts that learning should be strongest near the reward, where the teaching signal is largest. Over the first five outbound traverses in the Mask A maze we observed the opposite: turn error was lower at the holes *farthest* from the reward (Fig. S13E), which are precisely the holes the animal had already visited during its unrewarded sorties. In the Mask D maze, backward propagation predicts that preference should build up along the shortest path as a graded function of distance from the reward. Instead, the preference was concentrated at the bottleneck (Fig. 3E,F), and from off-path corridors the choice was near uniform for most start corridors with no trend across rewards (Fig. 3G,H). On the other hand, Endotaxis accounts for each of these: for the parallel learning of corridors and turns on the path-graph mazes (Fig. S13D,G), for the bottleneck choice and lack of preference among control corridors (Fig. 6C,D), and for the structural learning that precedes the first reward (Fig. 6F, Fig. S13A). In short, instead of acquiring competence over reward values, algorithms that extract latent structures may be the more promising route to few-shot learning and sample efficiency.

### 3.3 Maze learning by acortical mice

We found that mutant mice lacking a hippocampus and neocortex can nonetheless learn a complex maze. They took 2–3× longer to discover a rewarded route (Fig. 4), largely because of repetitive scanning across territory they had already covered. After just a few rewards, however, they traversed the maze as rapidly as normal mice (Figs. 4 and 5) and retained a functional memory of the experience over several weeks without exposure (Fig. 5). Every one of the five learning markers quantified above in wildtype mice was present in these animals. It is worth reviewing what is known about the mutants, and then asking how a brain without cortex or hippocampus can support such learning.

Studies of neural development have led to the creation of at least five strains of “acortical” mice [28, 41–44]. These constructs all follow the same principle: the Emx1-Cre driver line is used to introduce mutations in a gene essential for cell proliferation. Since Emx1-Cre is first expressed in the precursor cells of the dorsal forebrain [45], the Cre-induced mutation stops the development of that lineage. As a result, these strains share the same anatomical features: absence of hippocampal structures; absence of the dorsal cortex, with the remaining cortical volume reduced to 30% of normal; and absence of a corpus callosum (Figs. S6 and S7 and [28, 41–44]). The subcortical brain areas appear normal in size and arrangement (Figs. S6 and S7 and [41, 46, 47]).

Surprisingly, the mutants are quite proficient in many sensory-motor functions, including visual, auditory, olfactory, and somatosensory sensitivity [42, 46]; ultrasonic vocalizations [42, 48]; and motor control on the rotarod [28, 42]. They also succeed in simple learning paradigms, such as visual fear conditioning and a visually cued water maze [46]. While the mutant retains these basic functions, close inspection sometimes reveals quantitative differences from normal mice [46, 49].

At the circuit level, the absence of the sensory and motor neocortex also forces changes in the mutant’s subcortical connections. For example, the dorsal lateral geniculate nucleus of the thalamus (dLGN), which normally projects to the primary visual cortex, instead sends axons to some extracortical areas, and not to the remaining cortical tissue [47]. Similarly, neither the dLGN nor the superior colliculus (SC), which normally receive strong feedback from the primary visual cortex, receives any input from the remaining cortical tissue (Fig. S7) [47]. There are also second-order effects that do not involve cortical neurons directly: the projection from the retina to dLGN, though not to SC, is mostly absent in the mutant [46]. The acortical brain is therefore not simply a normal brain with the cortex removed, a point we return to below.

The present results extend the repertoire of acortical mouse behavior to include spatial learning in a complex maze and long-term memory of maze knowledge (Figs. 4 and 5). Read together, these patterns indicate that what fails in the mutant is limited to pre-reward exploration and latent learning (Fig. S8I). This outcome is surprising, because the hippocampus supports spatial navigation in rodents [14, 50], and the interaction between hippocampus and cortex is thought to establish long-term memories [51]. More specifically, the rodent hippocampus has been deemed essential for “place learning”, in which the animal uses an internal cognitive map, but not for tasks that are solved by learning a sequence of decisions, or by following visible beacons.

Several lines of evidence indicate that the Manhattan Maze challenges place learning specifically. The experiments were conducted in complete darkness, so the mouse has no access to external visual cues for orientation, nor to beacons that could signal the goal location from a distance. Further, landmarks within the maze are not the sole cue supporting navigation (Section 6.5). In a related study, we determined that normal mice do not rely on olfactory cues for spatial learning or navigation in these environments [13]. By elimination, the mouse appears to have available only the local shape of the corridor, as sensed by touch and whiskers, along with self-motion signals. Further, the maze solution does not require a memorized sequence. Efficient first traverses on the new path-graph mazes are rule-based rather than sequence-based (Fig. 5D-F, Fig. S11E). In the Mask D maze, the animal rarely executed the same turn sequence twice, all the while dramatically improving its performance (Fig. S12D-F). Both sensory evidence and maze design indicate that the acortical mouse operates using a cognitive map — the very representation the hippocampus is thought to build and store [14, 20, 52, 53] — and a reusable rule — the very function that is conventionally attributed to prefrontal cortex [17, 54, 55].

Conservatively, then, one can conclude that in the acortical mutant the remaining subcortical brain areas are capable of sustaining all of these interesting behaviors: few-shot learning of a navigation rule, credit assignment to decisions remote from a reward, and long-term memory of the maze environment. None of these functions strictly requires the special architecture and functionality of the six-layered cortex, or the trisynaptic circuit of the hippocampus, or the interaction between those areas.

Is this simply another instance of developmental compensation by which the brain responds to injury or defect [56, 57]? As discussed above, the mutant brain must be substantially rewired, given that neocortex and hippocampus are both source and target of so many neural projections. Any such rewiring, however, cannot have been driven by the exigencies of maze learning. Our animals were raised in a standard mouse cage, with no need or opportunity for long-range navigation. They were evaluated on a few hours of experience following their very first encounter with the maze environment. There was thus no opportunity for any task-driven compensation to create custom circuits of the type found after animals are over-trained on the same task for tens of thousands of trials [58].

Further, it is important to recognize that the mutant brain has some pronounced limitations. The failure of acortical mice to shelter during the visual looming response [59] likely follows from the absence of retrosplenial cortex, which has been implicated in the recognition and memory of a shelter [60]. Their inability to reach toward and manipulate fine objects with the forepaw (our unpublished observations and [28]) may result from the absence of a corticospinal projection to the digits, which is required for fine grasping skills [61]. In these cases, the behavioral defects are not graded but categorical, and each can be ascribed to a specific missing cortical circuit: the mutants exhibit the same lack of compensation one finds when lesioning or silencing the cortex in adult animals [59, 60, 62, 63]: Some functions simply do not recover after the lesion, and those indicate a strict cortical dependency.

By contrast, the navigation behavior of mutant animals is largely normal, excluding the initial exploration phase. While lesions or synaptic impairment of the hippocampus in adult animals have a clear effect on spatial learning [50, 64], they tend to delay the learning process, not abolish it. In the classic water maze experiments, the modified animals wasted time by circling inefficiently through the same search area, much as the acortical mice did here, and eventually they collected rewards and improved their performance [64, 65]. A parsimonious integration of the present results with prior lesion studies is that even in the normal mouse, a system entirely outside the hippocampus and neocortex can rapidly learn from exploration and rewards, and use the resulting spatial knowledge to guide subsequent navigation.

### 3.4 Limitations and open questions

The Manhattan Maze’s attractive attributes lie in the ability to generate a near-infinite diversity of environments, with graph structures far beyond what is possible in a 2-dimensional labyrinth. Furthermore, one can switch environments within seconds by replacing the mask. The Maze’s 3-dimensional structure also entails its chief limitation: the two stacked layers form a fully enclosed environment (Fig. 1A), which precludes wired neural recording. For now, the Manhattan Maze serves as a tool for fast behavioral screening in diverse spatial environments. The phenomena of spatial cognition identified this way can then be carried into open-top versions compatible with imaging, or paired with wireless recording, to combine behavioral and neural measurement.

Many interesting questions remain. For one, the sensory basis of these rapid learning phenomena is unresolved. We studied navigation in complete darkness and showed that it does not depend critically on landmarks the mice may deposit in the maze. Yet, how these spatial representations are acquired remains unclear, especially in acortical mice, which lack all primary sensory and motor cortices except the piriform cortex. Isolating individual sensory modalities alongside neural recording would determine which inputs are necessary for building cognitive maps.

Another enticing future direction is to compare brain processing in acortical and normal mice. The present study shows that hippocampus and neocortex are not strictly required for many aspects of spatial cognition. In all likelihood those brain areas do have a more subtle role in steering or coordinating the process. To resolve those contributions, one will want to silence hippocampus or cortical areas in a targeted and reversible manner during different stages of spatial learning and recall.

Finally, neural recordings in the acortical mutant could reveal candidate mechanisms for spatial cognition in the subcortical brain. Of particular interest are the sensory-motor transformations in the superior colliculus, the basal ganglia, and the cerebellum. Following the historical approach to the neuroscience of navigation, one could start with a survey of place cells, which seem to occur prominently in subcortical areas even in the wildtype mouse [66, 67].

## Supporting information

Vidoe V1

## 4 Acknowledgments

We thank Zeyu Jing and Jiang Wu for feedback on the study. This work was supported by the Simons Collaboration on the Global Brain (SCGB 543015 and 543025) to M.M. and P.P., and by NIH grant R01 NS111477 to M.M.

### 4.1 Author contributions

Conceptualization, J.Z., R.G., P.P., and M.M.; Methodology, J.Z., R.G., Z.T., P.P., and M.M.; Software, J.Z., R.G., and M.M.; Validation, J.Z. and M.M.; Formal Analysis, J.Z., R.G., J.Y.H., K.S., A.D., Z.T., and M.M.; Investigation, J.Z., R.G., J.Y.H., K.S., A.D., and Z.T.; Resources, M.M. and P.P.; Writing – Original Draft, J.Z. and M.M.; Writing – Review & Editing, J.Z., M.M., P.P., R.G., and Z.T.; Visualization, J.Z., R.G., and Z.T.; Supervision, P.P. and M.M.; Project Administration, J.Z., P.P. and M.M.; Funding Acquisition, P.P. and M.M.

## 5 Methods and Materials

### 5.1 Resource availability

#### 5.1.1 Lead contact

Requests for further information and resources should be directed to and will be fulfilled by the lead contact, Jieyu Zheng.

#### 5.1.2 Data and code availability

Processed trajectory data and analysis code are available at https://github.com/Jieyusz/Zheng_2026_Manhattan. Code for the Endotaxis Model is from https://github.com/markusmeister/Endotaxis-2023.

Any additional information required to reanalyze the data reported in this paper is available from the lead contact upon request.

### 5.2 Usage of artificial intelligence

During the preparation of this manuscript, we used Claude Code (Anthropic) to optimize the structure, readability, and efficiency of our custom data-analysis pipeline, and to format mathematical expressions into LaTeX syntax. The underlying scientific algorithms, data processing steps, and core mathematical formulations were entirely conceived, implemented, and validated by the authors. Additionally, ChatGPT-4o (OpenAI) and Claude Opus 4.8 (Anthropic) were used to assist with copy-editing, refining text clarity, and manuscript drafting. All AI-assisted and AI-generated outputs were reviewed and edited by the authors. We take full responsibility for the writing of the manuscript, the accuracy of the code, and the overall scientific integrity of the study. These tools were not used to generate original data, alter scientific interpretations, or influence the intellectual content of the work.

### 5.3 Experimental model and subject details

We used 47 adult C57BL/6J mice (Jackson Laboratory Strain 000664), 18 acortical mice, and 9 sibling mice (Section 5.3.1), totaling 33 male and 41 female mice, and analyzed 208 sessions from these 74 animals. The mice were aged between 61 and 106 days (mean 79.4 days) on the first day of the experiment. All were experimentally naïve and transferred to the same facility at least seven days before the experiments. The housing facility operated under a reversed light-dark cycle (13:11 Light:dark), and the experiments started and ended within 1 h of the dark cycle. The mice were transported in their cages from the facility to designated behavior test rooms on the experiment days. All procedures followed Protocol IA21-1656 and were approved by Caltech Institutional Animal Care and Use Committee (IACUC).

No statistical method was used to predetermine sample size: wildtype cohorts accumulated across mask configurations as the protocol developed, and the numbers of acortical and sibling mice were set by the breeding yield of the mutant colony (Section 5.3.1). Wildtype mice were assigned to mask-sequence groups by Latin square, so that littermates fell into different groups (Section 5.5.5). Genotype could not be randomized and investigators were not blinded to it, but no reported measure involves human scoring. All quantities come from automated tracking and scripted analysis (Section 5.6.2). None of the covariates — sex, starting age, experimenter, or test room — had significant effect on the learning measures (Section 5.6.3) after correction for multiple comparisons. The normal reference for acortical mice pool protocol-matched wildtype and sibling cohorts (Masks O, A, B and C) into a single control group. In the Mask D maze, they are reported separately.

#### 5.3.1 The acortical mice

We used a mutant mouse model which fails to develop its neocortex and hippocampus during embryonic development [28]. A transgenic mouse with Cre recombinase under the control of the Emx1 promoter was used to delete a conditional allele of Pals1, a critical gene involved in neuron proliferation. As a result, the entire lineage derived from these neuronal precursor cells is missing, including all projection neurons from neocortex and hippocampus. All the acortical mutant animals resulted from the conditional knockout of both copies of the Pals1 gene (genotype *Pals1*^loxp/loxp^:*Emx1-Cre*^+^, called “CKO” in Kim et al. [28]). The *Emx1-Cre*^+^ line was purchased from Jackson laboratory (Strain 005628) and the *Pals1*^loxp/loxp^ line was a gift from Seonhee Kim and Christopher Walsh [28]. We refer to the homozygous mutants in our experiments as the “acortical” mice, and their Cre*^−^* siblings as the “sibling” mice, to differentiate from the C57BL/6J “wildtype” mice in the previous sections. The genotypes of the acortical and sibling mice were confirmed by a commercial sequencing service (Transnetyx) using probes targeting *Emx1-Cre* and the *Pals1* loxP sites, and the brain phenotypes were verified by post mortem dissection to confirm the absence of neocortex and hippocampus.

To assess the conservation of subcortical structures, we first examined the midbrain regions involved in visual processing and sensorimotor transformation. The retinal ganglion cells send direct projections to the superficial layers of superior colliculus (SC). Using 3D gradient-echo Magnetic Resonance Imaging (MRI) sequences (Fig. S7A-B and Fig. S6A-D), we reconstructed the SC and periaqueductal gray (PAG) of an acortical mouse (Fig. S7C-E). These regions remain morphometrically comparable to those reported in wildtype animals, whereas the neocortex and hippocampus are entirely missing in the structural scans (Fig. S7B). These results confirm that acortical mutants retain intact midbrain structures while lacking both neocortex and hippocampus.

We next characterized the ventrolateral tissue remaining in the acortical mice. Anatomically, its location and structural characteristics resemble the allocortex rather than the six-layered neocortex. Next, we checked for connectivity between the SC and the piriform cortex. In wildtype animals, primary visual cortex (V1) sends direct projections to the superficial layers of SC. Using the retrogradely transported long-term non-transsynaptic virus, HSV-hEF1-mCherry, we confirmed these projections in the wildtype animal (Fig. S7F-G). In contrast, virus injections into acortical mice produced no retrograde labeling. We did not observe any projection neurons from the remaining 3-layer piriform areas (Fig. S7H-I). These results suggest that SC in the acortical mice receives no cortical input.

#### 5.3.2 Water deprivation

The mice were single-housed and water-deprived 20-22 h before the start of the experiment. For multi-day experiments, if a mouse obtained more water rewards than 25 ml/kg × body weight in the maze, it was returned to the housing facility without any water supply, continuing water deprivation until the next-day experiment. During the experiments and in the facility, the mice had ad libitum access to regular laboratory chow. Those that failed to obtain enough water rewards in the maze were supplemented with extra water to meet the minimal water requirement each day. The mice were also weighed at the beginning and the end of each day’s experiment to monitor weight loss due to water deprivation.

### 5.4 The Manhattan Maze

The Manhattan Maze is a grid-like three-dimensional labyrinth that can be easily reconfigured (Fig. 1A). The maze is constructed as two layers such that corridors in the maze are stacked on top of each other and separated by a mask in the middle (Fig. 1A). Each hole on the mask generates an intersection at its coordinate between two perpendicular corridors. We consider that a navigation decision is made when the mouse passes an open intersection: it may choose to move along the same corridor or to go through the hole, making a left or right turn into the perpendicular corridor (Fig. S1C). A series of holes on a mask creates a unique maze configuration or map, and a simple swap of the masks changes to a different maze configuration. Information about the physical maze construction can be found in the supplementary materials (Section 6.3).

#### 5.4.1 Mask design

Mask design for the Manhattan Maze was based on the graph representations at the corridor level. Theoretically, each mask is rendered as a graph with the corridors as vertices and the holes as edges. A two-layered 11×11 Manhattan Maze creates bipartite graphs: the eleven corridors in one layer form an independent set of disjoint vertices. Each corridor may have any number of holes from 0 to 11 where each hole gives access to any single corridor on the opposite layer. The number of all possible hole locations is 11 × 11, and the number of all possible combinations of these holes is 2^11×11^ ≈ 10^36^.

The middle corridors in the two layers have access holes leading to the water ports (Fig. 1A, see more in Section 5.5.1). The training Mask O (Fig. S1B) creates the shortest possible path by directly connecting these two corridors through the center hole. This mask also has the smallest size and order (P_2_) in this Manhattan Maze.

For the two-day learning task, we chose P_10_ (ten corridors and nine holes) masks based on a series of design constraints at the tile level (Fig. 1C). To reduce the bias of movement in different layers, we controlled the distances between holes in the vertical versus horizontal directions and the number of left turns versus right turns. After selecting Mask A, we picked Mask B to share the same turn sequence as A, but deliberately changed the hole locations so that the distances between holes were as different as possible (Fig. 2A). Mask C was chosen so that it had four different turns from the sequence of A (Fig. 2A). Together, Mask A, B, and C are isomorphic P_10_ graphs — i.e., sharing the same graph representation (Fig. 1C left) but using different subsets of corridors. These P_10_ masks were selected because most mice in the pilot experiments were able to learn them in a few hours.

Masks A, B, and C are termed “acyclic” because one cannot walk around and come back to the starting corridors without revisiting a corridor. A cyclic graph, however, can trap a wall-following agent in infinite walks in a subsection of the maze. Mask D is cyclic, specifically a union of two 4 × 4 bicliques (K_4,4_) and two path graphs: K_4,4_ + P_3_ + K_4,4_ + P_2_ (Fig. 3A). A biclique is defined as a graph with two sets of vertices where every vertex from one set is connected to all vertices from the other set. In each K_4,4_, a set of four corridors on one layer are all-to-all connected to another set of four corridors on the other layer. We designed the bicliques so that the corridors were roughly equally and symmetrically spaced. The bicliques are connected by short path graphs, serving as “bottlenecks.” To traverse the maze using the least number of corridors, a mouse must choose the correct hole to the corridor that is directly connected to the bottleneck path graphs.

For the intermediate stage of training in acortical mice, we designed two additional path-graph Masks E and F (details and results in supplementary materials). These two P_4_ masks have four corridors connected by three holes, which creates a three-turn sequence during traverse. Mask F is the center-symmetric version of Mask E.

### 5.5 Behavioral experiment

Throughout the experiments, the mice were handled individually and with minimal human contact. All animals were isolated from their cagemates at least one day before the experiments. Individual mice were transported and tested in separate testing rooms without any contact with other animals.

The mice started the experiment within the custom-made two-chamber home cage that was directly connected to the maze (Fig. S1A). The large chamber of the home cage contained bedding and food pellets on the floor. The small chamber served as a connecting passage between the large chamber and the maze, which reduced the amount of bedding dragged into the maze due to movement. Before each day of experiment, the maze apparatus and masks were thoroughly cleaned.

Before the experiment began, the home cage was disconnected from the maze. The mouse was put into the home cage and was allowed to acclimate to the chambers for at least fifteen minutes. Then the home cage was gently moved to align with the bottom tray of the maze, which allowed the mouse to freely explore the whole assembly. We define a *session* as the entire period where the mouse was allowed to enter and explore the maze freely while the mask configuration remains unchanged. Between sessions, the mouse stayed in the home cage that was disconnected from the maze, so that the experimenter could make changes by switching masks. At the end of each day, the mouse was trapped in and picked up from the home cage and then transported back to the animal facility for overnight rest.

#### 5.5.1 Reward delivery

The water rewards were delivered through the Sanworks Bpod State Machine r2.s controlled by a MATLAB script. The two water ports were placed close to the exit of the top and the bottom maze tray: one located at the end of the central vertical corridor in the top layer (the “Out” port), the other in the smaller chamber of the two-chamber home cage (the “Home” port). When the mouse pokes its nose in the port, that breaks an infrared beam and the port delivers a fixed-size drop of water. At the delivery of a reward, the infrared LED corresponding to the water port produces a flash and marks the reward events on video recordings.

At the beginning of each session, the mouse must first trigger the Out port, which delivers 10 µL water. Repeated poking at the same port no longer generates water rewards unless the mouse returns to the home cage to trigger the Home port. Upon poking, the Home port delivers 10 µL water and resets the Out port. Likewise, the Home port delivers water drops only once, until the mouse goes back into the maze and triggers the Out port. The reward delivery program motivates the mouse to alternate between the two water ports. The program was restarted every time the mask was swapped and a new session began.

#### 5.5.2 Training and handling

After several pilot experiments, we settled on two experiments for the wildtype mice: 1) a two-day protocol consisting of the same first day in learning Mask A and different second days for Masks A, B, and C, and 2) a one-day protocol with a different mask to learn (Mask D). Both experiments began with Mask O, which was necessary to teach the mice about the reward contingencies and acclimate them to the maze environments.

All mice went through the same training protocols in the Mask O maze (Fig. S1A-D). Most wildtype animals learned on their own and were able to alternate between the two water ports after a few hours. Should a mouse fail to trigger either port in more than 0.5 h, we gave a pulse to the port to generate a drop of water. In most cases, the mouse was motivated to lick the ports and subsequently learned the reward contingencies. On rare occasions where the mouse was unwilling to crawl through the hole, we guided the mouse through the center hole. The training on Mask O completed when at least twenty rewards are obtained at each port. Then the mouse was trapped in the home cage and changes were made to the maze. In any subsequent configurations, no additional instructions were provided to the wildtype mice. The mice were left to explore the mazes under the default automated reward delivery system without human interference.

#### 5.5.3 Training acortical mice

The acortical and sibling mice went through a similar training scheme, except for the assistance in the Mask O and Mask E mazes. After learning Mask O, a few acortical mice had difficulty learning to crawl through the second hole in a new Mask configuration. These mice received assistance. For example, a plastic petri dish under a hole served as a ladder for the mouse to crawl down, or the mouse was dropped down through the hole. For those circling in the home cage, they were trapped in the maze for increased exposure. With training, all acortical mice were able to learn Mask O. An intermediate Mask E maze (Section 6.4) was given to those that experienced difficulty crawling through more than one hole.

Tables 1 and 2 summarize the learning results of the acortical mice. Successful learners are defined as those that were able to obtain at least 20 rewards in a single session (maximum duration 8 h).

The long-term memory test of Mask A followed the general procedure as the overnight experiment in Section 5.5.2. During the breaks, the mice stayed in their home cages and had ad libitum food and water, until 20-22 h prior to the experiment when they were water deprived and separated. During the gaps, the mice had no access to Mask A. Because maze knowledge transfers across masks (Fig. 5D-F), we also audited whether any other mask was presented within these intervals. Of the 44 post-gap Mask-A sessions in Fig. 5B,C, 39 followed an interval containing no maze session of any kind. The five exceptions involve two acortical mice: mouse 683 was tested on Masks B, C and D on single days within three intervals (2, 6 and 22 days), and mouse 073 on Masks D, E and F within two intervals (2 and 7 days).

#### 5.5.4 Video recording

All videos were recorded during the dark phase of the light cycle (within 1-h difference) of the mice. The top cameras were located 1-2 meters above the arena and were controlled via FlyCapture and SpinView interfaces (Teledyne FLIR). The cameras captured both the home cage and the maze during the experiments (Fig. S1A). We also placed a webcam (Logitech) on the side of the maze for simultaneous recording, but the videos were not used for tracking or processing. All cameras recorded videos at thirty frames per second.

During the experiments, the entire arena was illuminated by 850-nm infrared LED lights that were invisible to the mice. During the mask swaps, the experimenters operated under red headlamps (wavelengths 610 - 760 nm) that temporarily illuminated the arena, while the mice were trapped in the home cage as the masks were changed. Mouse behaviors during human presence were recorded in the videos but not included in the analysis unless the mice entered the maze voluntarily before the experimenters left the testing rooms. In this case, the session time points were manually adjusted to include voluntary exploratory behaviors.

#### 5.5.5 Two-day testing of the path graph masks

For the path graph experiments, all mice continued to learn Mask A on the first day after completing the training in the Mask O maze. Only those that obtained at least 20 rewards in the Mask A maze moved on to the experiments on the next day. These animals had ad libitum access to food but were water deprived overnight. We only included data for those who completed the training on Day 1.

After completing the training stage, 25 mice progressed to the experiments on Day 2 where two novel masks, B and C, were introduced. We divided the full-day experiment into four 2-3-h sessions, each named after the mask used in that session following the order of “XYXZ” (X, Y, and Z represent three different masks). The mice were assigned to groups using Latin square to separate littermates into different groups, yielding groups of 3–6 mice. One mouse completed only the first three sessions, and one received a flipped Mask A in its second session; per-session sample sizes are therefore 24–25 (Fig. 2B).

We ended each session using performance-and time-based criteria, which allowed us to allocate sufficient time to all four sessions on Day 2 while accommodating variation in individual mouse performance. Session length was determined as follows: (1) If a mouse obtained more than 40 rewards on a repeated mask or more than 60 rewards on a new mask, we immediately switched to the next mask and began the next session. (2) If a mouse had spent 2 h on a repeated mask or 3 h on a new mask and had obtained at least 20 rewards, we proceeded to the next session. (3) If a mouse failed to obtain 20 rewards within this time window, we extended the session by an additional 0.5 to 1 h. These criteria yielded relatively balanced session durations while ensuring that mice had sufficient opportunity to perform in each maze.

### 5.6 Data analysis

#### 5.6.1 Animal tracking

We used DeepLabCut [68] Version 2.2rc3 and Version 3.0.0rc14 to track the location of the mouse in the maze. Each model was trained on 1000 frames from the pilot videos obtained in the same behavioral arena. To obtain reliable tracking results, we assigned twelve keypoints (Fig. S1A) to a mouse, including nose (1 keypoint), ears (2), head center (1), shoulders (3), body center (1), hips (3) and tail base (1). The trajectory data were processed based on the two-dimensional locations of the head center. Frames with low likelihood scores (i.e., frames in which the DeepLabCut model assigned a probability of less than 0.9 to the predicted keypoint location) were excluded, and missing keypoint positions were manually validated.

#### 5.6.2 Reduced trajectory data

We reduced the pixel locations of the head center into discrete tile coordinates in the maze assembly. The regions of interest of the entire maze assembly were manually drawn based on video screenshots. We define the location in the maze based on a unique set of (*x, y, z*) tile coordinates (Fig. S1B). The *x* and *y* coordinates of the mouse are determined by the head center keypoint tracked by DeepLabCut and its location in the grids. The *z* location is determined based on the time series. When the mouse crawls through a hole, only its *z* coordinate changes. We divide the time spent at the hole coordinate into two halves as an approximate for the time spent in the two tiles.

Next, the trajectories were segmented into *bouts* based on the maze coordinates. We pick the tiles adjacent to the water ports as the gates for segmentation. Since a mouse is typically two tiles long, we discard the bouts that are two tiles or shorter. Based on the starting and ending ports, we then categorize the bouts as H-H, H-O, O-H, O-O (H for “Home” and O for “Out”, see main text Section 2.1).

#### 5.6.3 Metrics of performance

To evaluate the performance of animals in the maze, we analyzed all bouts. The quality of bouts was assessed using temporal metrics, turn-based measures, and graph-level analyses.

First, the temporal metrics differentiated active navigation and goal-irrelevant time. We termed the entire duration of an experiment session as *session time*, which consisted of two parts: (1) time when the mouse was in the home cage or drinking at the ports (blank segments in Fig. 1D) and (2) *time in maze* when it was in the maze, including the time immobile in the maze (colored segments in Fig. 1D).

In the main text, we use time in maze to compute reward intervals, defined as the time between two rewards except for the first reward interval. Time to the first reward measures the duration of the first journey — i.e., from the beginning of the first bout to the end of the first traverse, when the first reward was obtained.

We then measured the durations of individual traverses. For curve fitting, the longest time per tile during traverse was capped to five seconds to remove sleeping and grooming episodes. The number of sorties between traverses was also used as a metric of efficiency.

Turn errors are scored at hole crossings, where the mouse commits to a direction on the corridor it enters. A turn is one out of two directional choices following a hole crossing (Fig. S1C). An error occurs when that direction deviates from the shortest path to reward (Fig. 1C and Fig. 2A). The turn error rate of a traverse is the count of incorrect first turns at each hole divided by the total number of holes. This binary and first-turn readout follows the turn-error measure used in Tolman’s T-maze [69, 70]: as turn decisions are treated as independent, the chance-level error rate is 0.5.

To evaluate performance in the context of each mask’s graph design, errors were quantified at two spatial resolutions: *corridor error* and *tile error*. An error is a step in the graph that increases the graph distance to the goal port. Tile error is sensitive to fine-scale decisions, including wrong turns at the holes, overshooting, and backtracking within the same corridor. A corridor error, in contrast, is generated only when the mouse crawls through a hole and enters a different corridor; turn direction at the holes does not affect it. Without crossing a hole, repeated scanning within the same corridor accumulates tile errors but no corridor errors. Both metrics are normalized to rates by dividing the raw error count by the total number of steps in that bout (corridor steps for corridor error, tile steps for tile error). To quantify within-corridor scanning directly, *tiles per corridor* is also reported, where a high value indicates more frequent scanning within the same corridors.

#### 5.6.4 Exponential curve fit

We used exponential curve fits to describe learning in traverse duration and turn error rate across repeated traverses. For each experimental condition, traverse duration was modeled as an exponential decay:

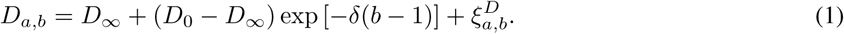

Here, *D_a,b_* denotes the duration of traverse *b* for animal *a*. The parameter *D*_0_ represents the fitted initial traverse duration, *D_∞_* represents the fitted asymptotic traverse duration, and *δ* is the non-negative learning rate. *D*_0_, *D_∞_*, and *δ* describe the learning curve of the population. Traverse index *b* is a positive integer, with *b* = 1 corresponding to the first traverse. The residual term 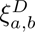 represents the deviation of an individual traverse duration from the fitted population-level curve. For nonlinear least-squares fitting, this corresponds to assuming mean-zero residual variability with constant variance, 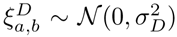, although statistical uncertainty in the fitted parameters was estimated by bootstrap rather than by the analytic covariance matrix. Similarly, turn error rate was modeled as:

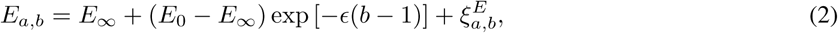

Here, *E_a,b_* denotes the turn error rate for animal *a* at traverse *b*. The parameter *E*_0_ represents the fitted initial turn error rate, *E_∞_* represents the fitted asymptotic turn error rate, and *ɛ* is the non-negative learning rate for turn-error reduction. For least-squares fitting, this corresponds to assuming mean-zero residual variability with constant variance, 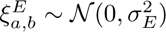.

Model parameters were estimated separately for each condition by ordinary nonlinear least-squares fitting (`scipy.optimize.curve_fit`). Rather than fitting animal-averaged summaries, the exponential model was fit directly to the pooled traverse-level observations from all animals in a condition, so that every individual traverse contributed to the fit. Statistical uncertainty was estimated with an animal-level cluster bootstrap. In each bootstrap iteration, animals were resampled with replacement within each condition, each resampled animal’s full traverse-by-traverse sequence was retained, and the model was refit to the resulting pooled dataset. The distribution of refit parameters across iterations defined the empirical bootstrap distribution and percentile confidence interval for each parameter.

For each fit, we constrained all parameters to the same predefined bounds and initialized the optimizer with fixed starting values. For traverse duration, the bounds are 2 ≤ *D_∞_* ≤ 60, 5 ≤ *D*_0_ ≤ 800, and 0.01 ≤ *δ* ≤ 1. For turn error rate, the bounds are 0.001 ≤ *E_∞_* ≤ 0.5, 0.1 ≤ *E*_0_ ≤ 1, and 0.01 ≤ *ɛ* ≤ 1. Split exponential fits between outbound and homebound traverses were examined, but they are only well-powered for the 25-mice Day 1 Mask A dataset (Fig. 1G), so a consistent pooled treatment is used throughout.

#### 5.6.5 Relative magnitude estimate

To evaluate performance in the Mask A maze after breaks (Fig. 5B-C and Fig. S15B), we quantify relative traverse duration and relative turn error rate. For each later Mask A session, the performance is compared to the first 10 traverses with the same mouse’s Day 1 baseline. Relative performance is calculated as the mean value in the later session divided by the mean value from the Day 1 baseline.

Group-level 95% confidence intervals are estimated using a hierarchical bootstrap with 1000 iterations. In each iteration, sessions are sampled with replacement. For each sampled session, the traverses are sampled with replacement from the first 10 traverses of that session and independently sampled traverses with replacement from the corresponding mouse’s Day 1 baseline. The mean for each resampled later session is then divided by the mean for its resampled Day 1 baseline to generate a relative performance value. These relative values are then averaged across sampled sessions. The 2.5th and 97.5th percentiles of the bootstrap distribution are used as the 95% confidence interval.

To quantify the magnitude of differences between fitted parameters, we calculated parameter ratios between conditions A and B using the bootstrap distributions from the curve fits. For a fitted parameter *θ*, such as *D_∞_*, *D*_0_, *δ*, *E_∞_*, *E*_0_, or *ɛ*, the ratio is computed for each bootstrap replicate:

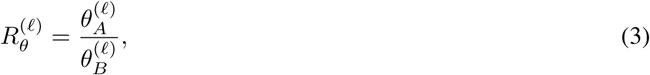

where 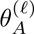 and 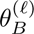 are the fitted parameter estimates from the *ℓ*th bootstrap replicate for the two conditions being compared. Thus, confidence intervals for change estimates are obtained directly from the bootstrap distribution of the ratio, rather than by dividing the confidence limits of the individual parameter estimates.

For Day 2 and Day 1 comparisons, the parameter ratios are computed for each fitted parameter *θ*. For each Day 2 session and mask condition, the ratio of the Day 2 parameter estimate to the corresponding Day 1 estimate is expressed as

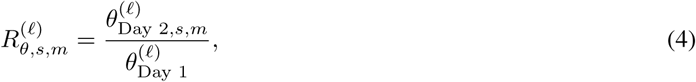

where *s* denotes the Day 2 session, *m* denotes the mask condition. To summarize the overall Day 2 effect across sessions and masks, we compute the median ratio across all Day 2 session–mask combinations within each bootstrap replicate:

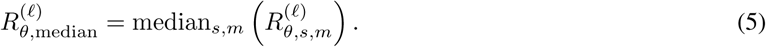

Initial-performance parameters, *D*_0_ and *E*_0_, are used directly because they are well identified by the fitted curves. For rate parameters *δ* and *ɛ*, bootstrap draws within a fractional tolerance of 10*^−^*^3^ of either fitting bound are treated as boundary-saturated and excluded from ratio calculations.

Because the fitted asymptotes *D_∞_* and *E_∞_* are often weakly identified, the late-performance estimate is used for ratio comparisons instead. Using *b*_late_, the latest traverse number observed in every session being compared, late performance is computed by evaluating the fitted exponential curve at *b*_late_:

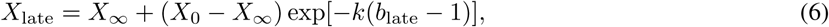

where *X* denotes either traverse duration *D* or turn error rate *E*, and *k* is the generic learning rate (*δ* for *D* and *ɛ* for *E*). Ratios are then computed from paired bootstrap iterations of these fitted late-performance values with the same *b*_late_, and 95% confidence intervals are taken from the percentile bootstrap distribution. These late-performance quantities are reported as *D*_late_ and *E*_late_.

#### 5.6.6 Directional transitions in the Mask D maze

To evaluate learning in the Mask D maze (Fig. 3E-H, Fig. 5H), we quantified directional transitions between corridors. Each bout is represented as a directed transition-count matrix describing movements between corridors. Each entry of the matrix corresponds to the number of times a transition from one corridor to another was used. Because transitions are directional, movements from corridor *c_i_* to corridor *c_j_* are counted separately from movements from corridor *c_j_* to corridor *c_i_*.

Thus for each bout *q*, the corridor sequence is converted into a directed transition-count matrix *T* ^(*q*)^, where each element 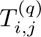 represents the number of times the animal moved from corridor *c_j_* to corridor *c_i_* during bout *q*. Throughout, matrix indices follow the convention that the row is the destination and the column is the source: each column of *T* ^(*q*)^ describes the outgoing transitions from one corridor, and each row describes the destination corridor. Fig. 3E–F focus on choices made after the animal reached the bottleneck junction. The transition counts are summed across all the bouts during a journey,

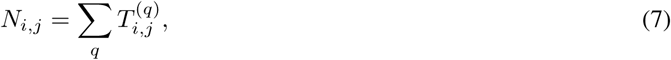

for calculating the *choice ratio* from the gateway corridor *c*_B_ to a possible next corridor *c_j_* as

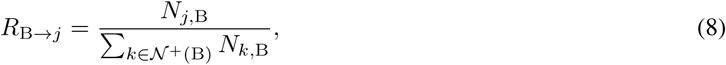

where 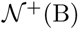 is the index set of a chosen group of target corridors that can be reached directly from the bottleneck junction, for example, all five neighbors of the gateway corridor in Fig. 3E-F and the three off-path corridors for each biclique corridor in Fig. 3G-H (see below). This ratio gives the fraction of all outgoing bottleneck transitions that were directed toward corridor *c_j_*. Transitions from *c*_B_ to corridors outside 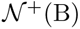 are excluded from this conditional ratio. Under a uniform-choice null restricted to the target set, the chance probability for each target corridor is 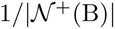.

We also quantified choices among corridors that did not lie on the shortest path (Fig. 3G,H). In the biclique region of Mask D, excluding the shortest-path corridors leaves an off-path subgraph with the structure of K_3,3_: three vertical corridors and three horizontal corridors remain available as off-path alternatives. Because each corridor had three equivalent off-path alternatives under this restricted analysis, the uniform-choice null expectation was 1*/*3 for each target.

#### 5.6.7 Traverse similarity in the Mask D maze

For Fig. S5, the similarity between traverses is based on the directed corridor transitions they used. Each traverse is represented as the set of directed transitions used at least once, ignoring how often each transition occurred. For two traverses, *T* ^(*p*)^ and *T* ^(*q*)^, an adjusted Jaccard similarity is computed:

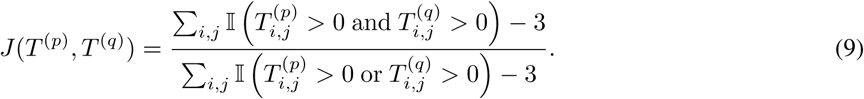

Here, 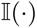 is the indicator function, which equals 1 when the condition inside the parentheses is true and 0 otherwise. We subtract three because these transitions through the bottlenecks were mandatory and shared by all valid traverses (Fig. 3A). Since each traverse uses at least five transitions, the denominator will not become zero. Thus, *J*(*T* ^(*p*)^*, T* ^(*q*)^) = 0 indicates that two traverses shared only the obligatory transitions, whereas *J*(*T* ^(*p*)^*, T* ^(*q*)^) = 1 indicates that they used the same set of directed transitions.

For each session with *m* outbound traverses and *n* homebound traverses, three similarity matrices are constructed: outbound-outbound, homebound-homebound, and outbound-homebound,

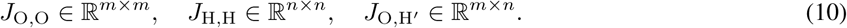

In each matrix, rows and columns correspond to individual traverses, and each entry gives the adjusted Jaccard similarity between the corresponding pair of traverses. Let 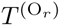 denote the transition matrix of the *r*-th outbound traverse, and let 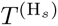 denote the transition matrix of the *s*-th homebound traverse. The within-direction similarity matrices are defined as and

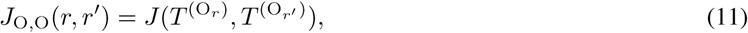

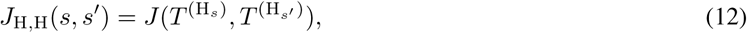

The outbound–homebound comparison uses the transposed transition matrix of homebound traverse to check if the mouse reused the same transition in reverse while returning. Specifically, for each homebound traverse H*_s_*, a reversed homebound traverse H*′_s_* has a transition matrix

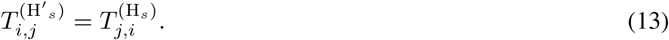

The outbound-homebound similarity matrix is then defined as

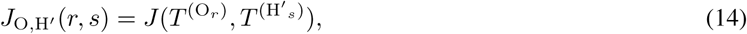

The diagonal elements of *J*_O,O_ and *J*_H,H_ are 1 by definition. To quantify how transition usage changed over time, we compute the mean similarity along the *k*-th off-diagonal of each within-direction matrix (Fig. S5C). This measure captures how similarity between traverses depends on their temporal separation.

### 5.7 Modeling

#### 5.7.1 First-order Markov model with forward bias

To compare animal exploration against non-learning baselines (Fig. S1I), we model a Markov walk on the maze graph and compute the expected number of steps required to travel from a designated start to a designated goal. This is described by a one-parameter family of first-order Markov models indexed by a *forward bias β* [71]: at *β* = 0.5 the walker is memoryless, choosing uniformly among all available moves with no dependence on the path taken to arrive there; for *β >* 0.5 it preferentially advances into new nodes. A node of the graph can be a corridor or a tile (Section 5.4.1).

At each step the walker occupies a directed edge (*j* → *i*), currently at node *i*, having arrived from node *j*; and moves to a neighbor *l* of *i* (*A_li_*= 1 in the adjacency matrix *A* of the maze graph) with probability

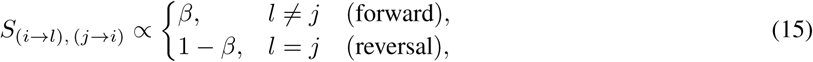

normalized over the neighbors of *i*; at a dead end (where *j* is the only neighbor) the walker is forced to reverse. The bias *β* ∈ (0, 1] sets the tendency to advance into new nodes rather than retreat, and, collected over all directed edges, these probabilities form a column-stochastic transition matrix *S*.

Indexing the directed-edge states by *u, v*, the completion time is the first-passage time of this chain to the goal node *e*, made absorbing by marking every state that steps into *e*. Writing *τ_u_* for the expected steps to first reach *e* from state *u*,

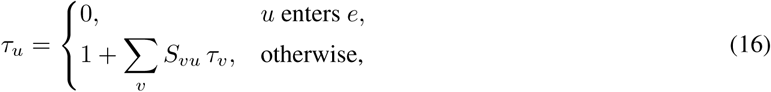

where the leading 1 counts the step just taken. Deleting the absorbing states leaves the transient vector ***τ*** *′* and substochastic matrix *S′*, giving

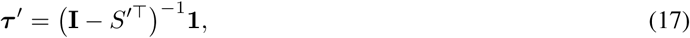

with fundamental matrix **F** = (**I** − *S^′⊤^*)*^−^*^1^ [72]. The completion time from a start node, *τ̅*(*β*), averages Eq. (17) over the edges leaving that node (the walker’s possible first moves) and adds 1 for that first step; it is finite as long as the start and end nodes are connected.

At *β* = 0.5 the forward and reversal probabilities are equal, so Eq. (15) becomes independent of the incoming node *j* and the chain reduces to a memoryless walk over nodes with *S_ij_*= *A_ij_/* ∑*_k_ A_kj_*; at *β* = 1 the walker never reverses.

The acyclic P_10_ graph, evaluated at the corridor level (Section 5.4.1 and Fig. 1C) from the Home to the Out corridor, gives the expected time of a memoryless walk *τ̅* (0.5) = 81, and of a perfect forward walk *τ̅* (1) = 9, also the shortest path (Table 3).

Cyclic Mask D (K_4,4_ + P_3_ + K_4,4_ + P_2_; Fig. 3A) gives *τ̅* (0.5) = 166.75 and *τ̅* (1) = 92.59, both far above its shortest path of 5, as the bicliques trap walkers with limited memory (Table 3).

The transition rule Eq. (15) yields a direct estimate of an animal’s forward bias from its own path. At an interior node of degree *g* = ∑*_l_ A_li_*, a walker arriving from *j* has one reversal move (weight 1 − *β*) and *g* − 1 forward moves (each weight *β*), so Eq. (15) assigns a reversal probability

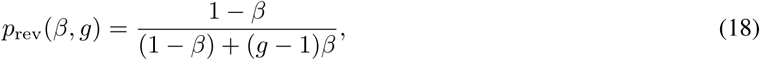

Writing *c*(*t*) for the corridor occupied at the *t*-th corridor decision, we count a reversal whenever the animal returns to the corridor it just left, *c*(*t* + 1) = *c*(*t* − 1). Over *N*_dec_ corridor decisions containing *N*_rev_ reversals, with *g*(*t*) = deg(*c*(*t*)) the degree at decision *t*, the empirical forward bias 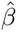 is the value that makes the model’s expected reversals match the observed count,

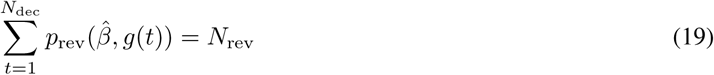

Further, the fundamental matrix gives the expected corridor errors — i.e., transitions that increase the graph distance to the goal (E(*β*), Section 5.6.3), which decreases monotonically with *β*. Because the corridor graph is bipartite, every step changes the goal distance by one, so the expected walk length is *τ̅* (*β*) = 2E(*β*)+*L*, where *L* is the shortest-path length in holes. Normalizing errors by walk length gives the per-step *corridor error rate*, 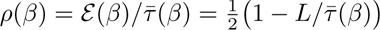, which approaches the memoryless chance level 0.5 as the walk lengthens. The memoryless walker (*β* = 0.5) has a corridor error rate of 0.44 in the P_10_ maze and 0.49 in the Mask D maze, dropping to 0 and 0.47 at *β* = 1 (Table 3).

The same reasoning carries over to tile error rate. The tile graph is also bipartite, so every step changes the tile-distance to the goal by exactly ±1, and the same calculation 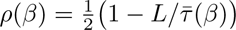 applies. At tile resolution the memoryless walker’s expected traverse length *τ̅* far exceeds the shortest-path length *L*, so the null is near 0.50 for every mask (A 0.496, B 0.496, C 0.496, D 0.498).

#### 5.7.2 The Endotaxis Model

We used the Endotaxis model [32], a neuromorphic algorithm that constructs a map through unrewarded exploration and uses local signals for navigation without requiring global shortest-path calculations, as a candidate algorithm for learning the corridor graphs.

Writing *b* for the traverse index (*b* = 1 the first traverse) and *d* for the distance to reward, the Mask A corridor error rate *ρ* and turn error rate *E* share the same predicted form at every genuine decision position,

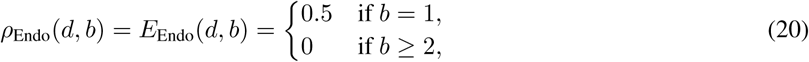

chance on the first traverse and 0 thereafter, independent of *d* (for the corridor metric the forced dead-end start corridor is 0 for all *b*; the turn metric has no such position).

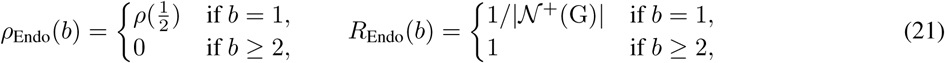

where 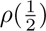 is the corridor error rate of the memoryless 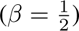 walker for that direction (0.49 outbound; Table 3), and G is the gateway corridor into the bottleneck, with |*N* ^+^(G)| = 5 so chance is 0.2. These are the analytic Endotaxis predictions plotted in Fig. S13D,G (Eq. (20)) and Fig. 6C,D (Eq. (21)).

For simulations of animal journeys, we used the first two journeys from the mouse shown in Fig. 3B as the input trajectory for Mask D (Fig. 6E-G) and the first two journeys from the mouse in Fig. 1E for Mask A (Fig. S13A). The model received only the sequence of corridor visits made by the animal and was not given the full corridor graph in advance. Each corridor was treated as a discrete location. As the animal moved through the maze, the model updated the map-cell connections according to the observed transitions between corridors. We then examined the learned map and the goal signal at selected time points before and after the first rewarded traverse.

We used the following learning parameters: gain *γ* = 0.21, learning threshold *θ* = 0.2, goal-learning rate *α* = 0.2, and decay = 0. These values were selected to demonstrate the operation of the model. The gain *γ* controls the propagation of activity through the learned map, the threshold *θ* determines when map-cell connections are updated, and *α* controls the update of goal-cell synapses following reward encounters [32]. The chosen values allow the goal signal to propagate through the learned map while maintaining sufficient contrast in node activities for visualization. Similar qualitative behavior was obtained over a broad range of parameter values (data not shown). We did not attempt to optimize these parameters to fit the behavioral data but to illustrate the qualitative operation of the model.

#### 5.7.3 Model-free Reinforcement Learning

We used a tabular, model-free Q-learning agent as the null reinforcement learning (RL) model, applied both to Mask D (Fig. 6) and to Mask A, the P_10_ linear path graph (Fig. S13). The two travel directions were treated as independent problems. On Mask D we read out the corridor error rate per traverse and the bottleneck choice ratio. On Mask A we read out the corridor error rate of each corridor and the turn error rate at each hole, ranked by distance to reward.

Learning follows the off-policy TD(0) update:

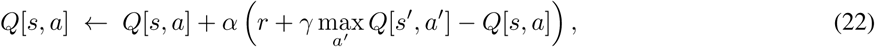

where *s* is the current state, *a* the action, *Q* the action-value table, *r* the reward, and *s′* the next state; max*_a_′ Q*[*s′, a′*] is the greedy bootstrap estimate of the next state’s value, *α* is the learning rate, and *γ* the discount factor. Reward is +1 on reaching the direction’s goal port and 0 otherwise.

For the corridor agent, states are corridors and actions are steps to an adjacent corridor through holes. On Mask D the same construction applies to the full corridor graph, where the gateway corridors afford five actions and the mandatory bottleneck two. On Mask A the nine corridors before the end corridor afford two actions at each interior position, and the start corridor is a forced single-neighbor dead end. For the turn agent, states are the 9 holes of Mask A and actions are allocentric headings *a* ∈ {*N, S, E, W*} (table *Q*[hole, dir]; Fig. S1C).

On Mask A the agent is trained on each animal’s sorties only: for every journey it first replays the leading sorties, and we read out the agent’s error on the traverse *before* that traverse’s reward updates *Q*, matching how the animals are scored. The traverse is then replayed with its reward to update *Q*. For turns we score the turn error rate as for the animals (Section 5.6.3), which leaves two candidate headings per hole, so chance is 0.5.

As a result of TD(0), reward value propagates back exactly one position per rewarded traverse. The single-direction Mask A learning curve has an exact closed form and the reported RL prediction is analytic. For a genuine decision position at distance-to-reward *d* on traverse *b*, the corridor error rate is

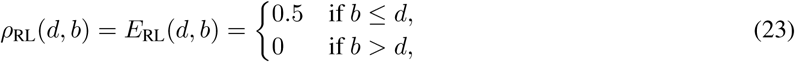

i.e. chance for the first *d* traverses and 0 thereafter (the error rate at the start corridor is 0 for all *b*). The staircase follows from three facts: sorties reach no reward and leave every *Q* at 0, so the agent is at chance at all positions on traverse 1; TD(0) advances the low-error frontier one position per rewarded traverse; and the greedy readout makes the per-position update independent of *α* or the random walk. This closed form was confirmed with trained per-animal simulations of both agents, aggregated over 25 animals per direction, which reproduced the staircase.

Mask D admits no such closed form. Its corridor graph joins two bicliques through a bottleneck corridor, so the corridor path is not unique and reward value must propagate across bicliques. The max operator in the TD update is nonlinear, and the rate at which value reaches the bottleneck depends on how the self-played traverses pass through the bicliques. As on Mask A, the pre-reward sorties carry no reward and leave *Q* unchanged, so each traverse was generated by self-play: a memoryless random walk to the goal, with the reward delivered and *Q* updated on arrival, and the readout taken before the reward. We report the mean over 20 independent agents, which sets the error bars in Fig. 6A-D.

## 6 Supplementary Materials

### 6.1 Supplementary figures

**Supplementary Figure S1:**
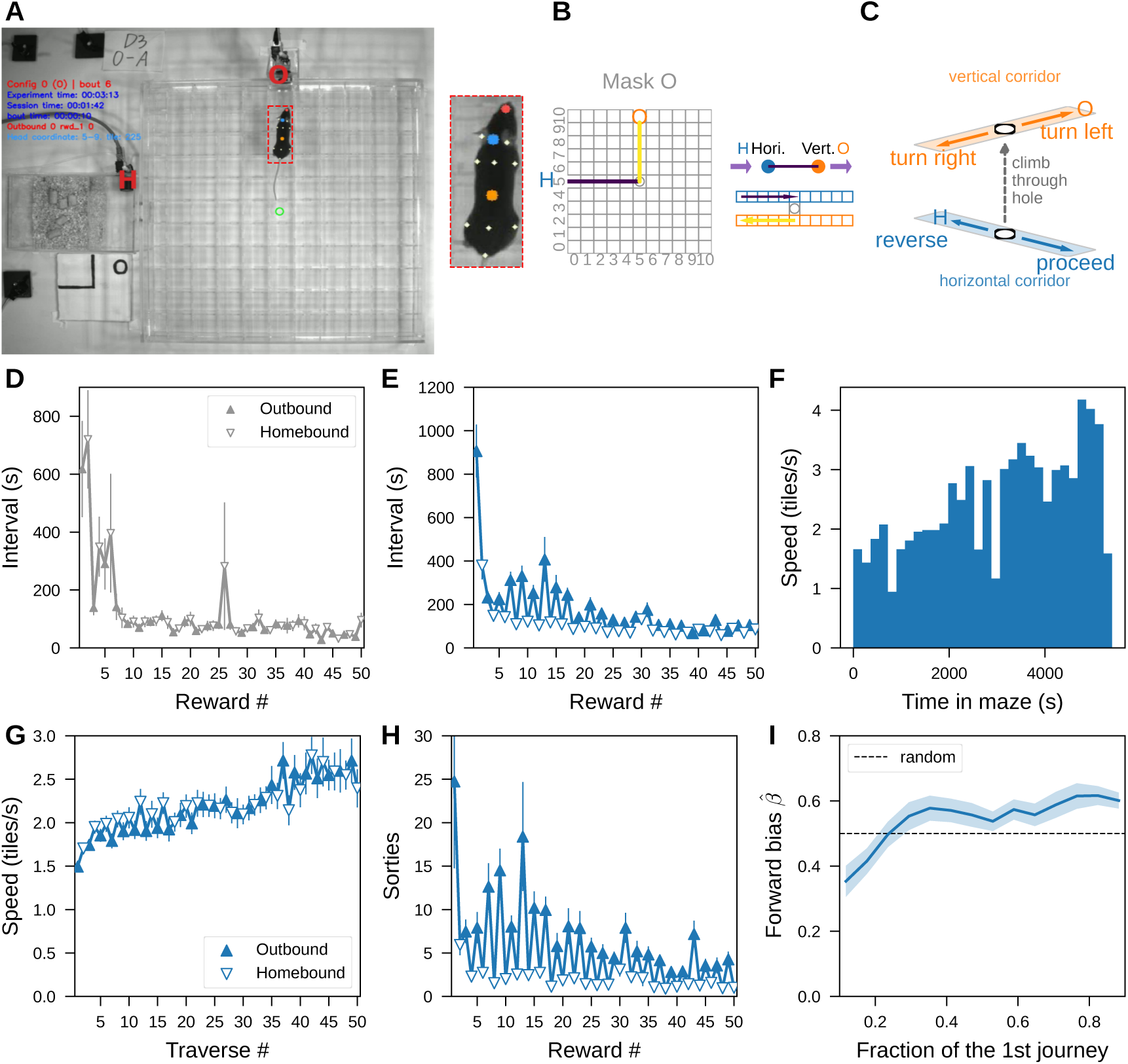
Supplementary results for Day 1 experiments. **A-D. Learning Mask O. A.** Top view of the arena with Mask O. A mouse cage is attached to the left entrance; it includes the Home water port (H). The Out water port (O) is connected to the top entrance. The mouse was tracked with 12 keypoints (right), with the head center (blue) used for trajectory segmentation (Section 5.6.2). **B.** Left: Top view of the shortest path in the Mask O maze. The tile coordinates are indexed from zero at the bottom left corner. Right: Graph representation of Mask O. Top: the graph of Mask O in corridors. Bottom: the hole segmented the corridors into a left turn from H to O. **C.** Four decisions can be made at a hole: from the horizontal (blue) corridor, the mouse can climb up the hole and make a left or right turn in the vertical corridor (orange arrows), or stay in the corridor to proceed or reverse from the hole (blue arrows). **D.** Reward intervals (in session time) on Mask O (25 mice) shortened quickly after the first two rewards. **E-I. Learning Mask A.**Lines and errorbars (or shade) show mean ± SE across animals. **E.** Reward intervals in the Mask A maze (25 mice). **F.** Speed in the Mask A maze by Wildtype 1 in Fig. 1D (bin size = 3 minutes and one tile is 1.5 inches). **G.** Traverse speed by all mice in Fig. 1G (25 mice). **H.** Number of sorties between rewards. **I.** Forward bias (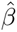, Section 5.7.1) of the mice before the first reward. Line smoothed with a moving average over 0.2 of the journey.

**Supplementary Figure S2:**
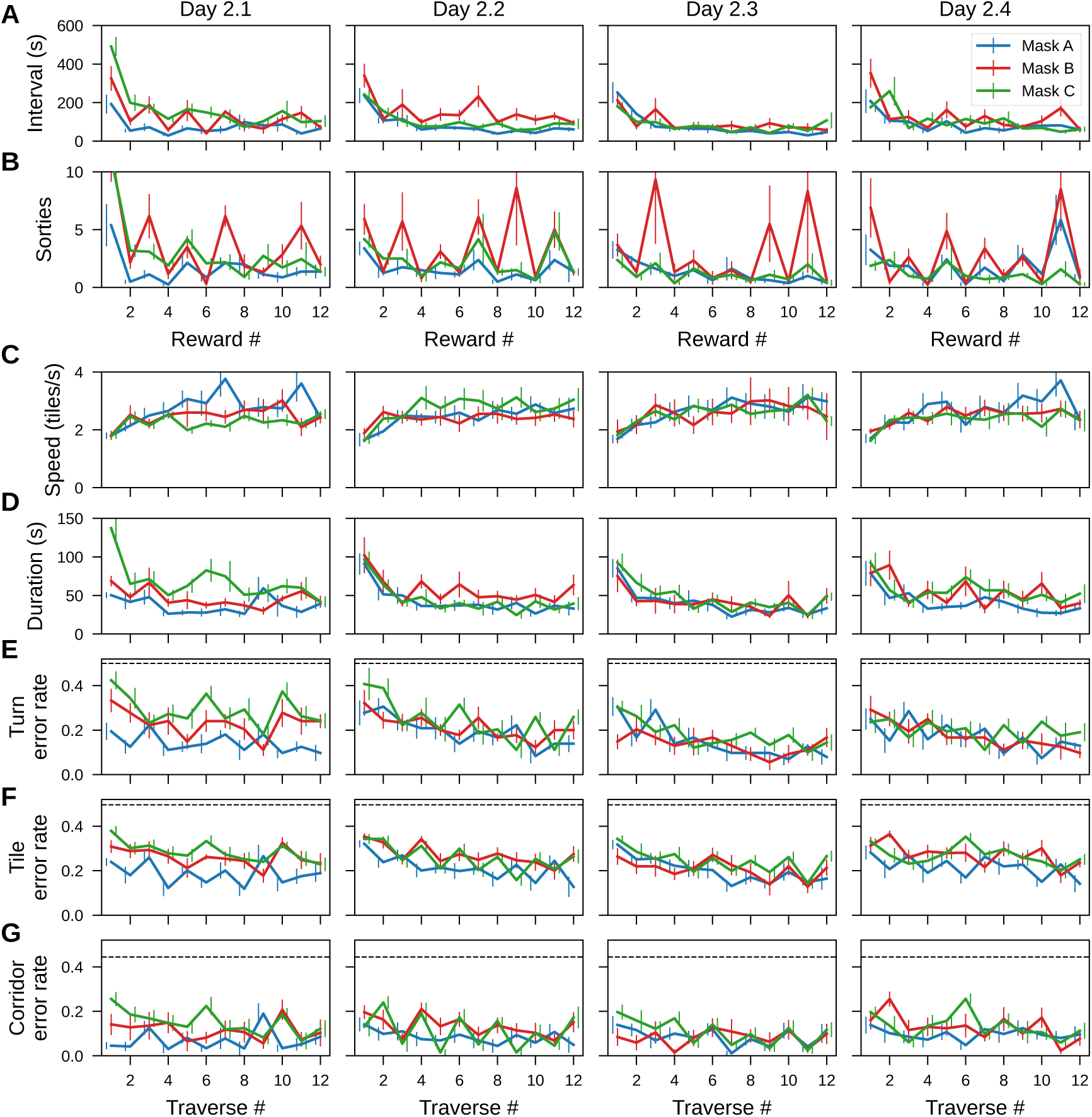
Additional metrics of performance on Day 2. Lines and errorbars show mean ± SE across animals. Dashed lines, chance error rates. **A.** Reward intervals across all four sessions, grouped by masks. **B.** Number of sorties between rewards. **C.** Traverse speed (tiles/s). **D.** Traverse duration. **E.** Turn error rates of traverses. **F.** Tile errors of traverses. **G.** Corridor errors of traverses.

**Supplementary Figure S3:**
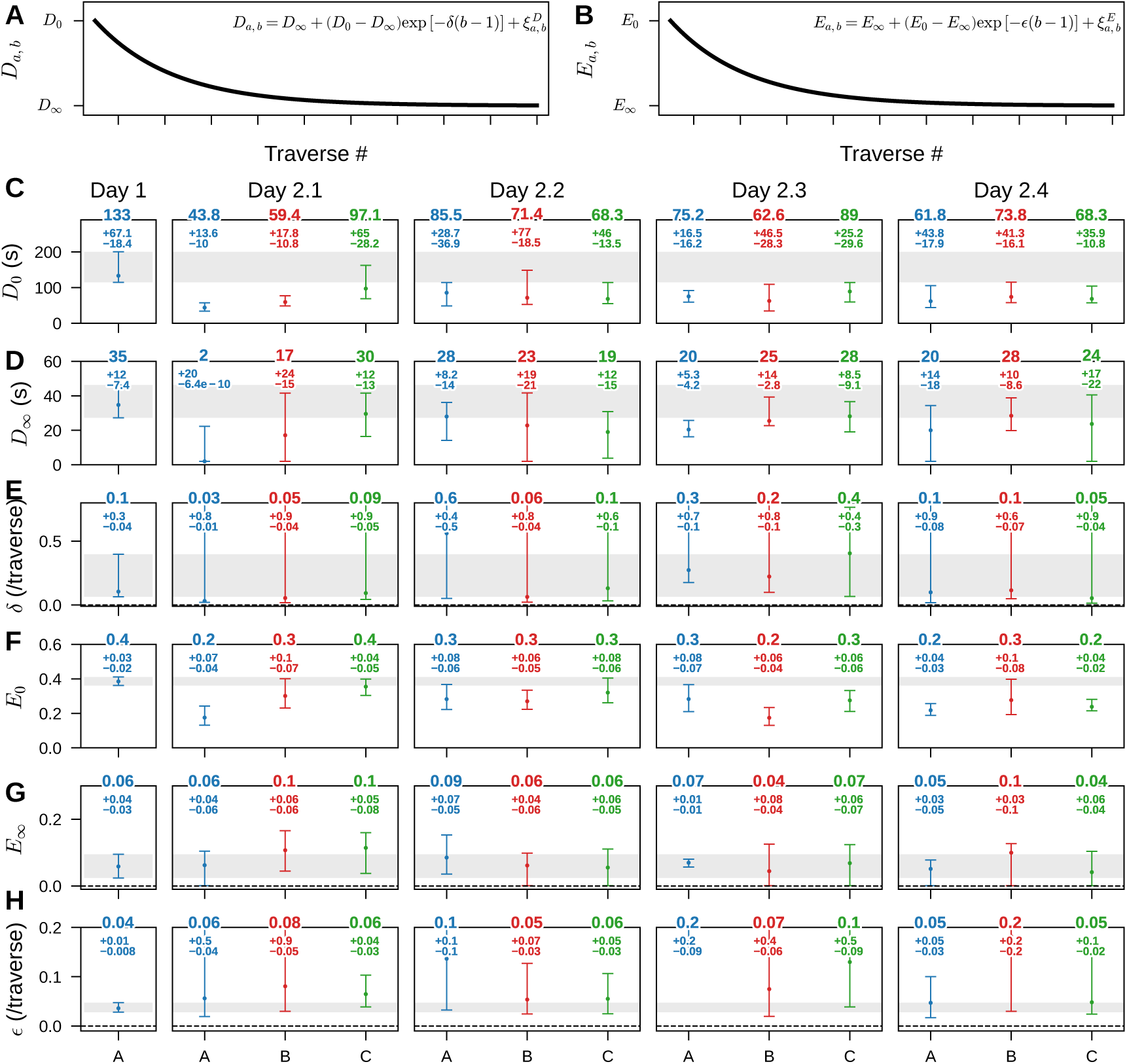
Confidence intervals of curve-fit parameters in the two-day experiments. **A-B.** Equations and schematics of the fitted learning curves (See Section 5.6.4). **C-H.** Mask parameters are differentiated by colors: Blue for Mask A, red for Mask B, and green for Mask C. Numbers denote the point estimate, with 95% confidence intervals shown as asymmetric upper and lower deviations. The gray shades show the 95% confidence intervals of the parameters from Day 1. **C-E.** Parameter fits for traverse duration. **C.** *D*_0_: duration of the first traverse in seconds. **D.***D_∞_*: asymptotic traverse duration in seconds. **E.** *δ*: learning rate. **F-H.** Parameter fits for traverse turn error rates. **F.** *E*_0_: turn error rate of the first traverse. **G.** *E_∞_*: asymptotic turn error rate. **H.** *ɛ*: learning rate.

**Supplementary Figure S4:**
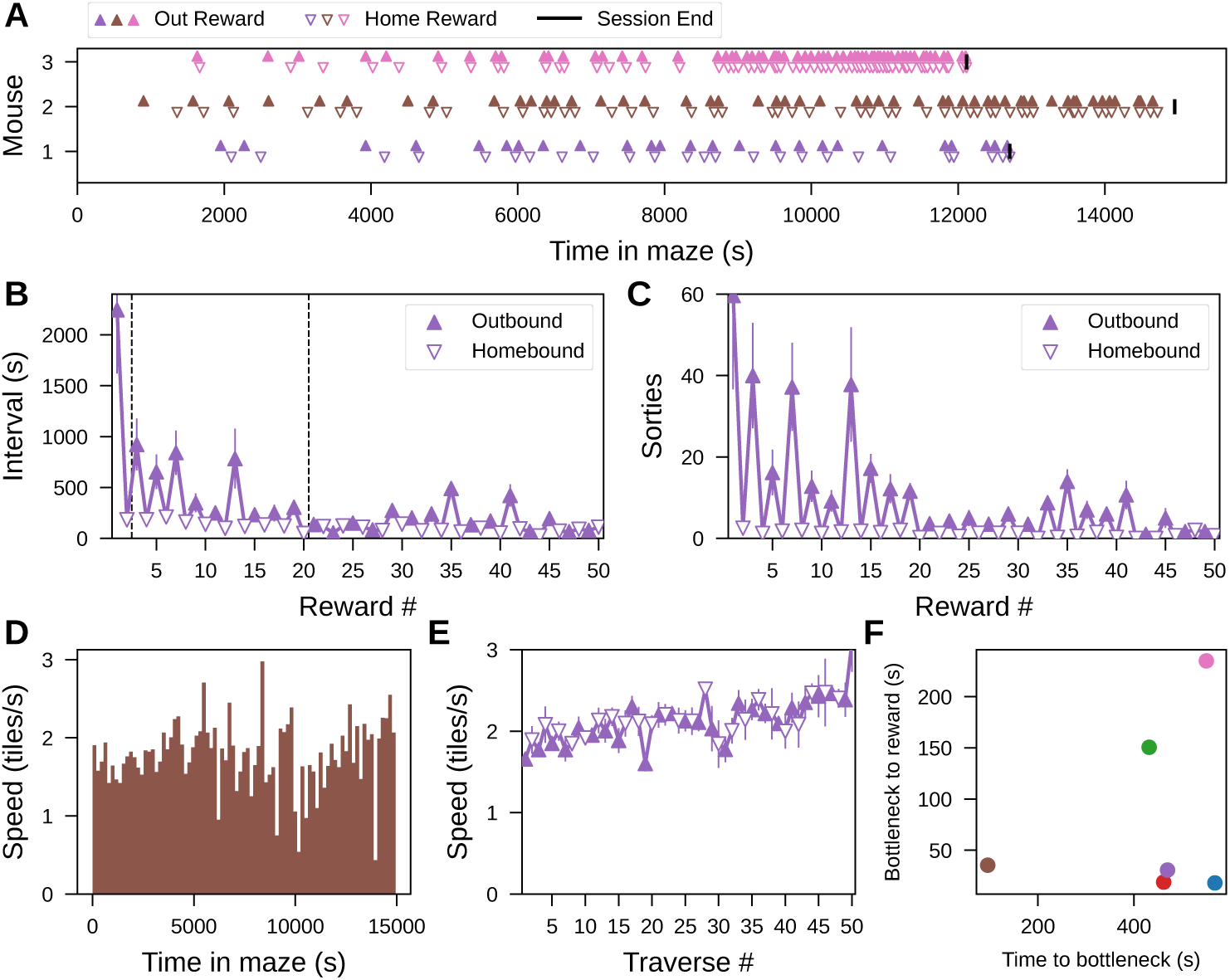
Additional metrics of learning Mask D as the first mask. **A.** Rewards obtained over time in maze by three mice in the Mask D maze. Black vertical bars mark the end of sessions. **B.** Reward intervals (6 mice, mean ± SE across animals). **C.** Number of sorties between rewards (mean ± SE). **D.** Speed of Wildtype 1 in Panel A during the entire session. **E.** Traverse speeds of all mice. **F.** Time (in maze) to reach the bottleneck for the first time vs. time (in maze) between the last visit of the bottleneck and the first reward. Points are individual animals.

**Supplementary Figure S5:**
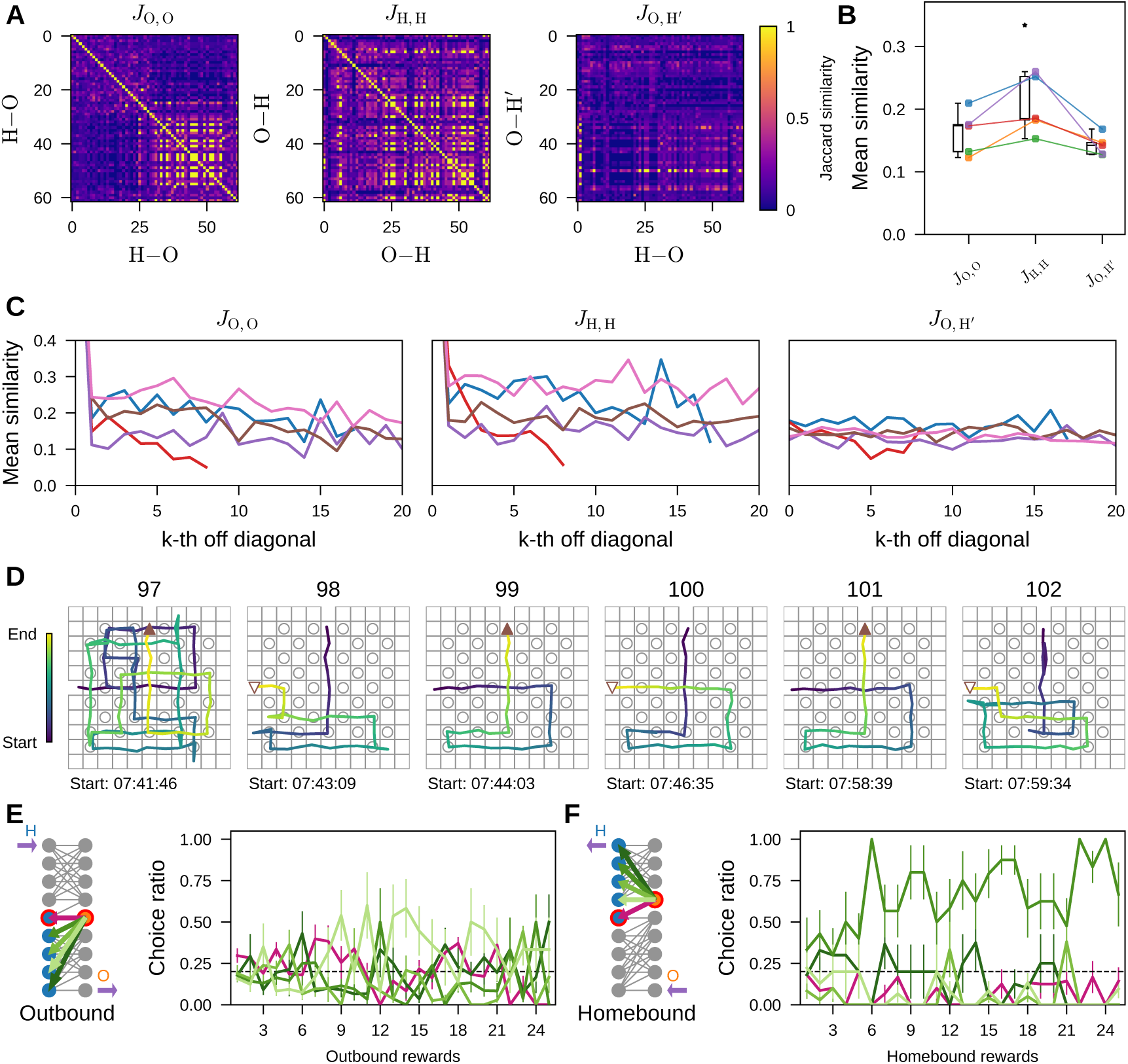
Varied routes in the Mask D maze. **A.** The Jaccard similarity (Section 5.6.7) between the transition vectors in bicliques in outbound traverses (*J*_O,O_), homebound traverses (*J*_H,H_), and traverse pairs (*J*_O,H_*′*) by the mouse in Fig. 3B. **B.** Mean similarity values off-diagonal for all three groups of vectors (5 mice). Each colored point is a unique animal (Friedman test, *χ*^2^(2) = 8.40, *p* = 0.015, paired Wilcoxon signed-rank test not significant). **C.** Mean similarities over the distance off diagonal (each colored line is a unique animal that corresponds to B). **D.** Temporally adjacent traverses by the mouse in Fig. 3B. **E-F.** At the symmetric point of the bottleneck, preference for reversing back to the bottleneck (pink) compared to chance (horizontal dashed line, 0.2) and the other corridors (green).

**Supplementary Figure S6:**
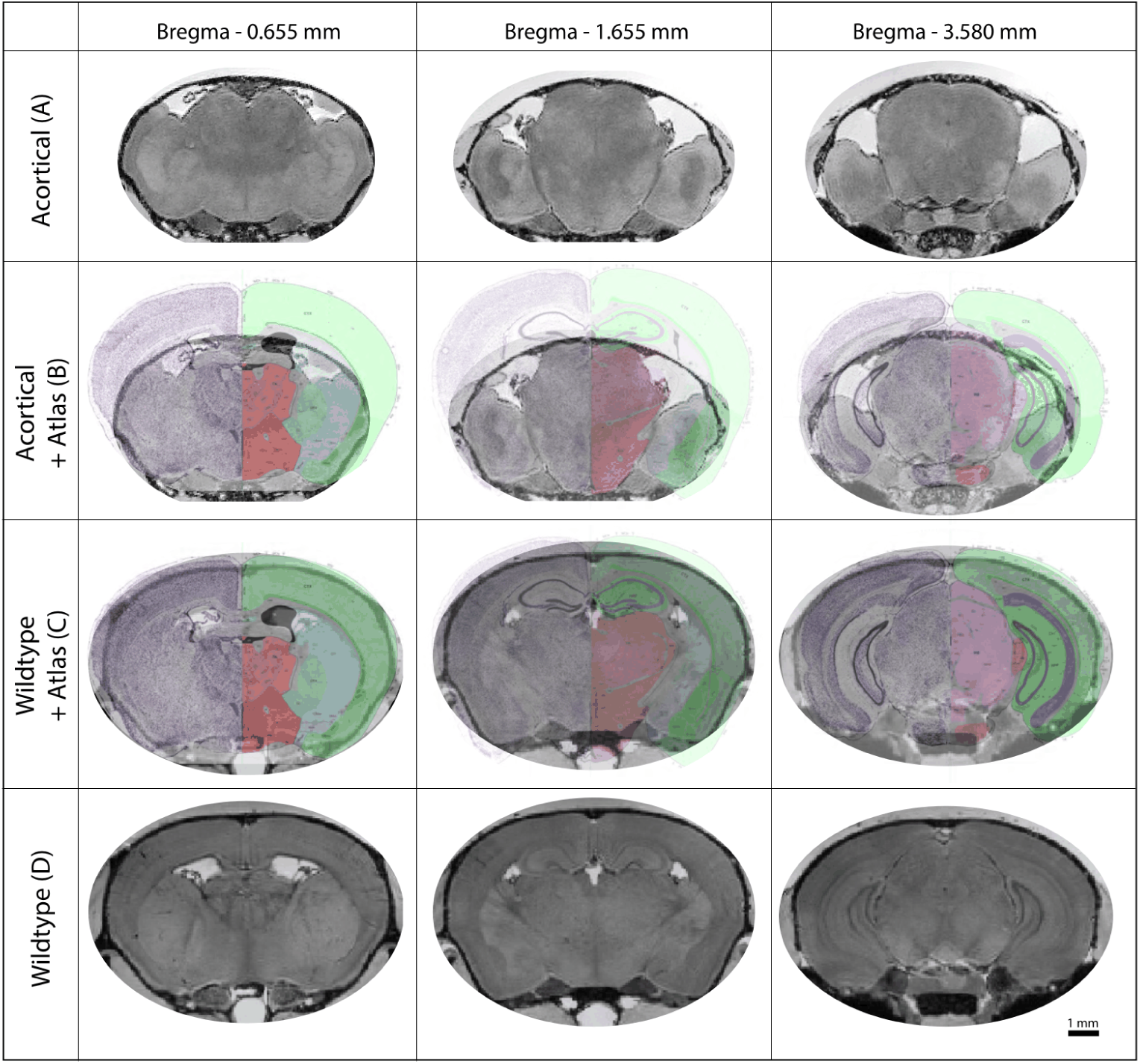
Mutant mouse with no neocortex and no hippocampus. Coronal MRI sections at three anterior-posterior locations (columns): Bregma −0.655 mm, Bregma −1.655 mm, and Bregma −3.580 mm. The corresponding coronal sections are overlaid with atlas rows from the Allen Institute Mouse Brain Reference Atlas [73, 74]. Scale bar, 1 mm. **A.** Sections from an acortical mouse. **B.** The same sections as A with the atlas overlay. **C.** Sections from a wildtype mouse with the atlas overlay. **D.** The same wildtype sections without overlay.

**Supplementary Figure S7:**
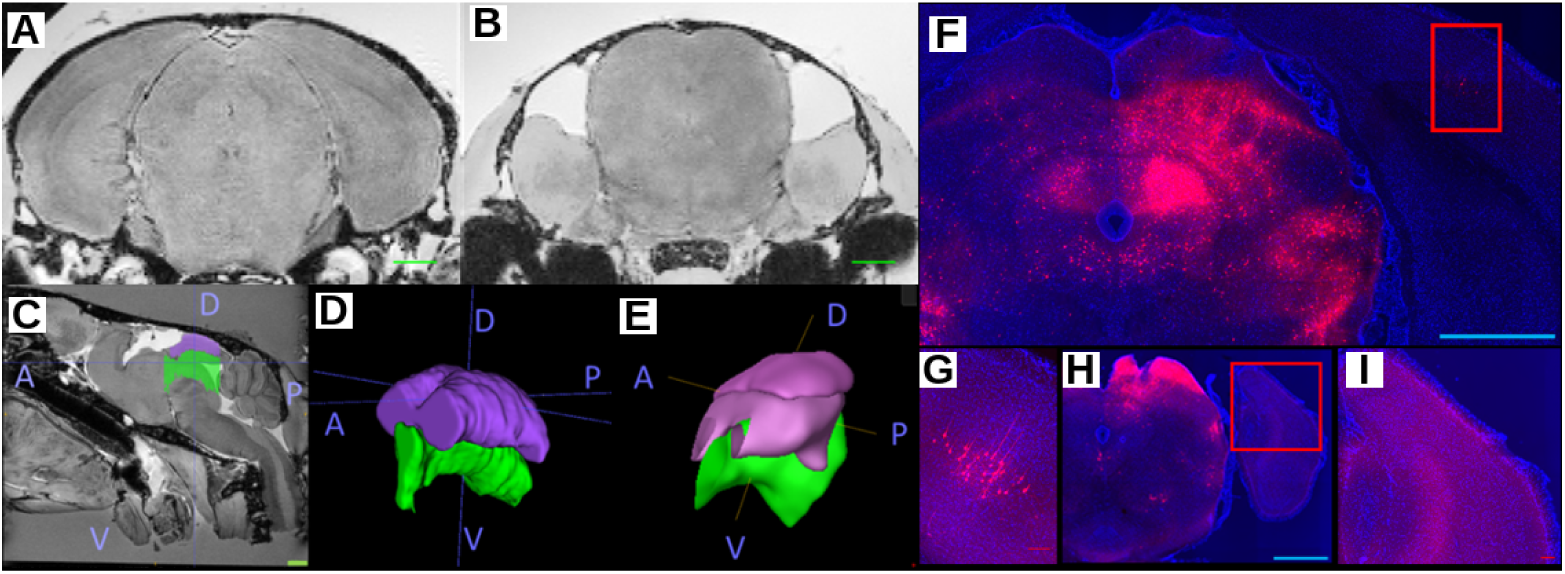
Midbrain structures of the mutant mouse. Superior colliculus in the mutant mouse does not receive cortical input. **A-B.** High resolution T2*-weighted 3D gradient echo images of wildtype (A) and mutant (B) mice. **C.** Sagittal section of mutant mice showing superior colliculus (SC) in purple and periaqueductal gray (PAG) in green. Blue axes show anterior (A), posterior (P), dorsal (D) and ventral (V) directions. **D-E.** 3D reconstruction of SC and PAG in mutant (D) and wildtype mice (E); scale bar, 1 mm. **F-I.** Retrograde labeling of projection neurons in superficial SC via long term HSV-hEF1*α*-mCherry. SC in wildtype mice (F) receives projections from primary visual cortex (inset and G). Acortical mice (H) lack any cortical input (inset and I) to SC; scale bar, 1mm.

**Supplementary Figure S8:**
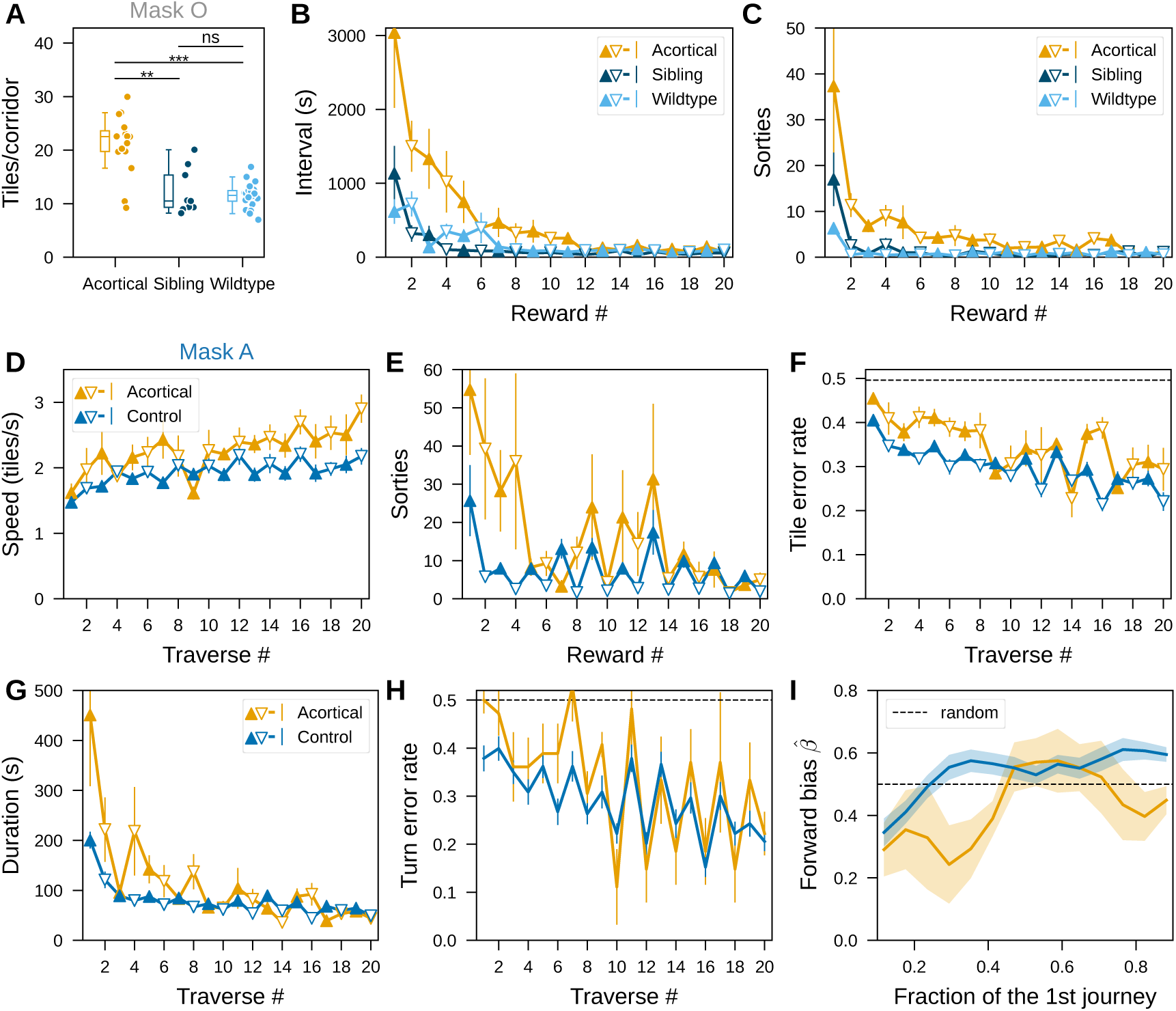
Slower initial learning of acortical mice. Metrics comparing acortical (orange), sibling (dark blue) and wildtype (light blue) mice. Lines and error bars show mean ± SE across animals. **A-C.** In the Mask O Maze. **A.** Tiles per corridor in the first two rewards compared across acortical (15 mice, orange), sibling (9 mice, dark blue) and wildtype (25 mice, light blue). Asterisks mark significant two-sided pairwise Mann-Whitney U tests (acortical vs. sibling, *U* = 119, *p* = 0.002; acortical vs. wildtype, *U* = 335, *p* = 4.0 × 10*^−^*^5^; sibling vs. wildtype, *U* = 105, *p* = 0.78, n.s.) after a significant Kruskal-Wallis test (*H* = 18.7, *p* = 8.8 × 10*^−^*^5^). **B.** Reward intervals (session time) of acortical, sibling and wildtype mice. The light blue curve is identical to Fig. S1D. **C.** Number of sorties. **D-I.** In the Mask A maze. Control mice (blue) are pooled sibling and wildtype mice. **D.** Speed of traverses. **E.** Number of sorties. **F.** Tile error rate of traverses. Dashed line, chance error rate. **G.** Traverse duration. **H.** Turn error rate of traverses. Dashed line, chance error rate. **I.** Forward bias (*β*^^^, Section 5.7.1) along the first journey (mean ± SE). The dashed line marks the memoryless walker (*β* = 0.5). Lines smoothed with a moving average over 0.2 of the journey.

**Supplementary Figure S9:**
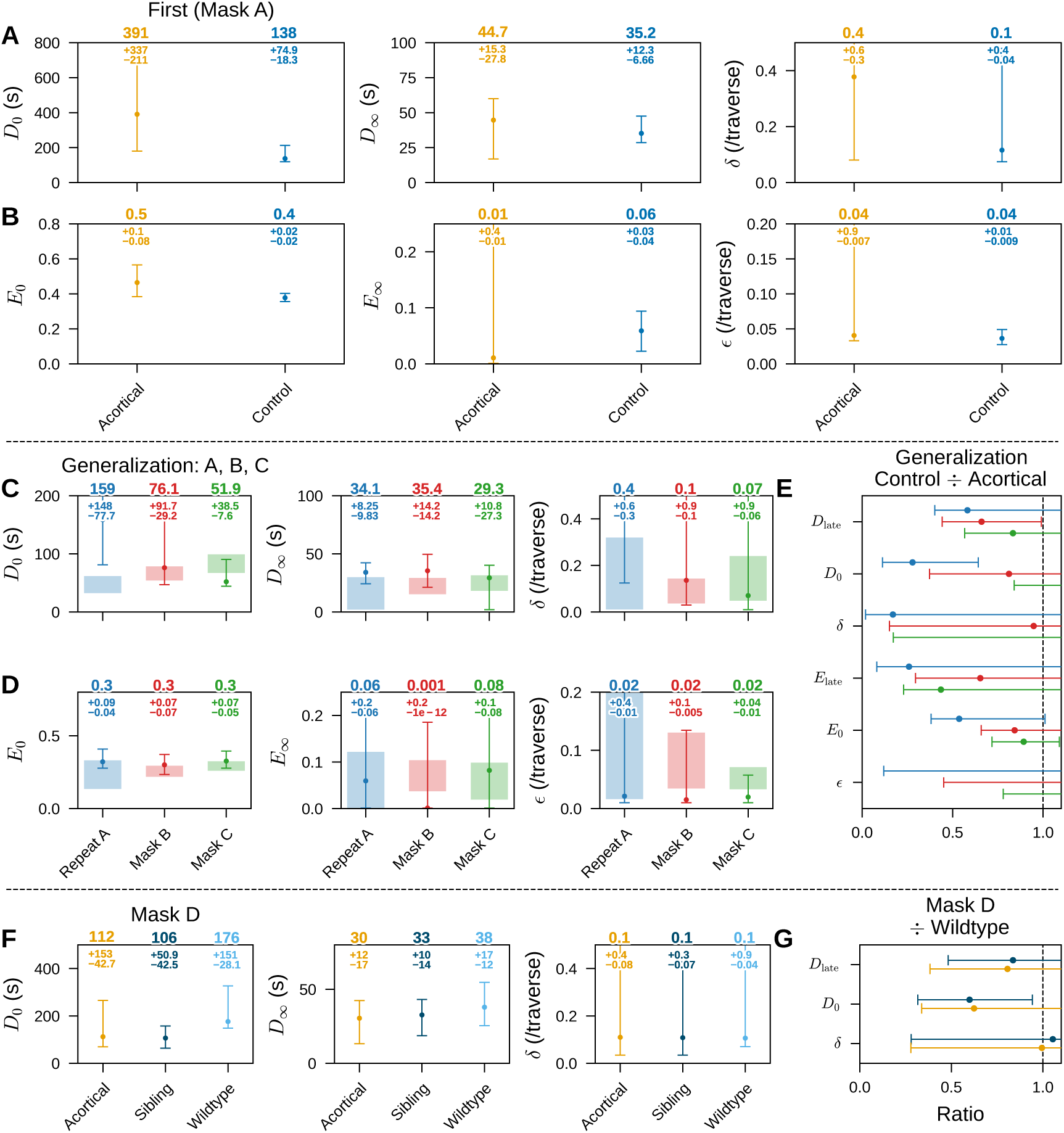
Curve fit results between acortical and control mice. Numbers denote the point estimate, with 95% confidence intervals shown as asymmetric upper and lower deviations. **A-B.** Learning Mask A as the first mask across acortical (4 mice, orange) and control (27 mice, blue). **A.** Parameter fits for traverse duration. *D*_0_: duration of the first traverse in seconds. *D_∞_* duration of the optimal traverse in seconds. *δ* learning rate. **B.** Parameter fits for traverse turn error rates. *E*_0_: turn error rate of the first traverse. *E_∞_* turn error rate of the optimal traverse. *ɛ* learning rate. **C-D.** Exponential fit parameters for traverse duration (C) and turn error rate (D) on new masks and repeated Mask A. The shaded regions plot the same parameter fits for the control group: pooled for Mask B (28 mice) and Mask C (31 mice), and sibling mice alone for the repeated Mask A (7 mice). **E.** Generalization parameter ratios for both traverse duration and turn error rate across the repeated Mask A, Mask B and Mask C, computed as control/acortical. Each marker is the bootstrap median ratio and its 95% percentile confidence interval (Section 5.6.5); a ratio of 1 (dashed line) indicates no difference between cohorts. **F.** Exponential fit parameters for traverse duration in the Mask D maze across acortical (5 mice, orange), sibling (8 mice, dark blue) and wildtype (6 mice, light blue). **G.** As in E, Mask D duration-parameter ratios relative to wildtype: sibling/wildtype (dark blue) and acortical/wildtype (orange).

**Supplementary Figure S10:**
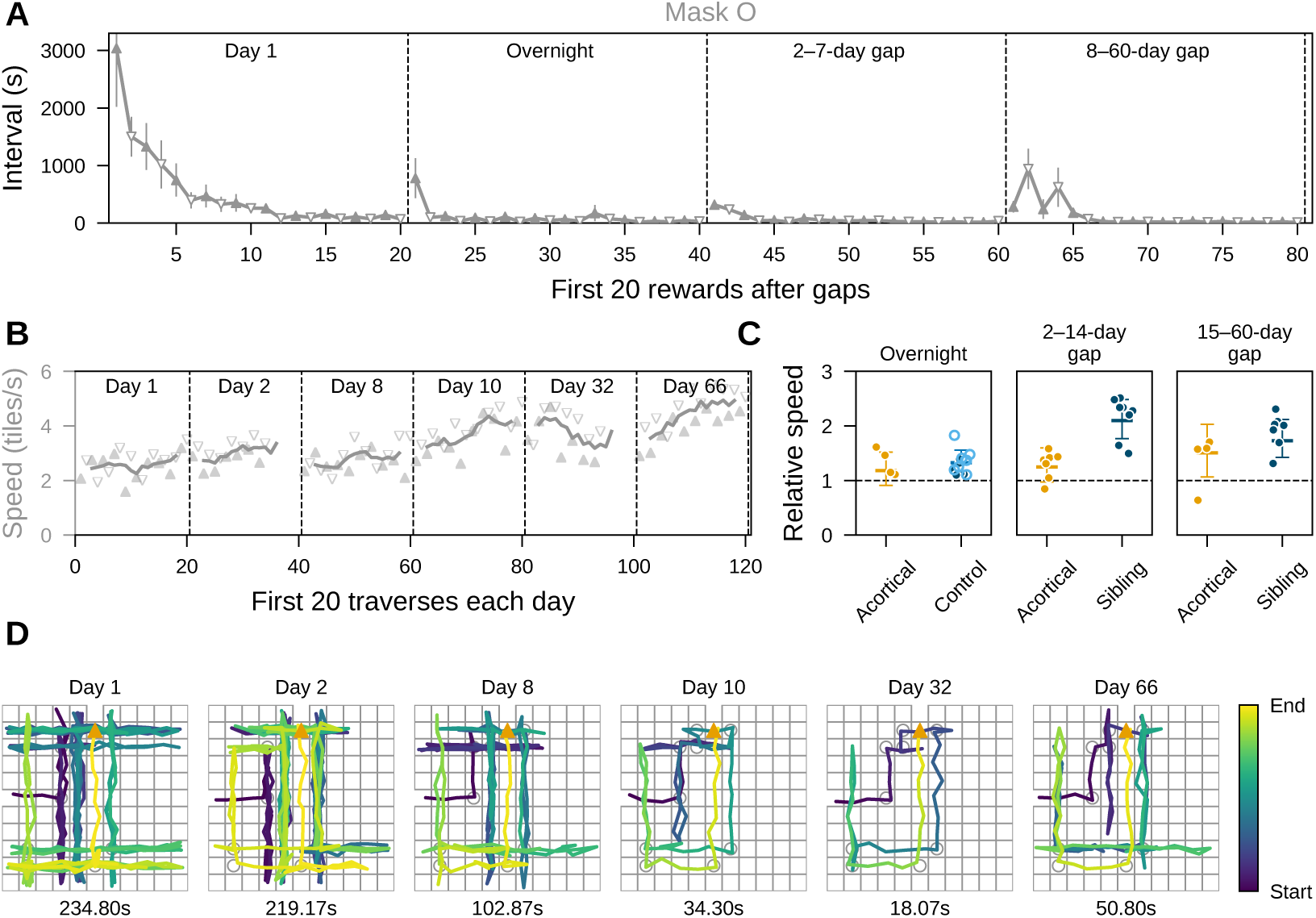
Long-term memory of masks. **A.** Memory of Mask O in acortical mice (14 mice). Reward intervals (session time) of acortical mice in their first session overnight and after 2-7 days or 8-60 days of gap. The first panel is identical to the orange curve in Fig. S8B. **B.** Traverse speed (tiles/s) of the mouse in Fig. 5A. Solid lines are moving average over five traverses. **C.** Relative mean traverse speed in acortical (6 mice, orange), sibling (8 mice, dark blue), and wildtype (8 mice, light blue). The error bar shows 95% confidence intervals of the population-level mean, calculated from the pooled traverses across all sessions within each gap range (see text). Dots are from individual sessions. **D.** First traverse on each day in the Mask A maze by mouse in Fig. 5A.

**Supplementary Figure S11:**
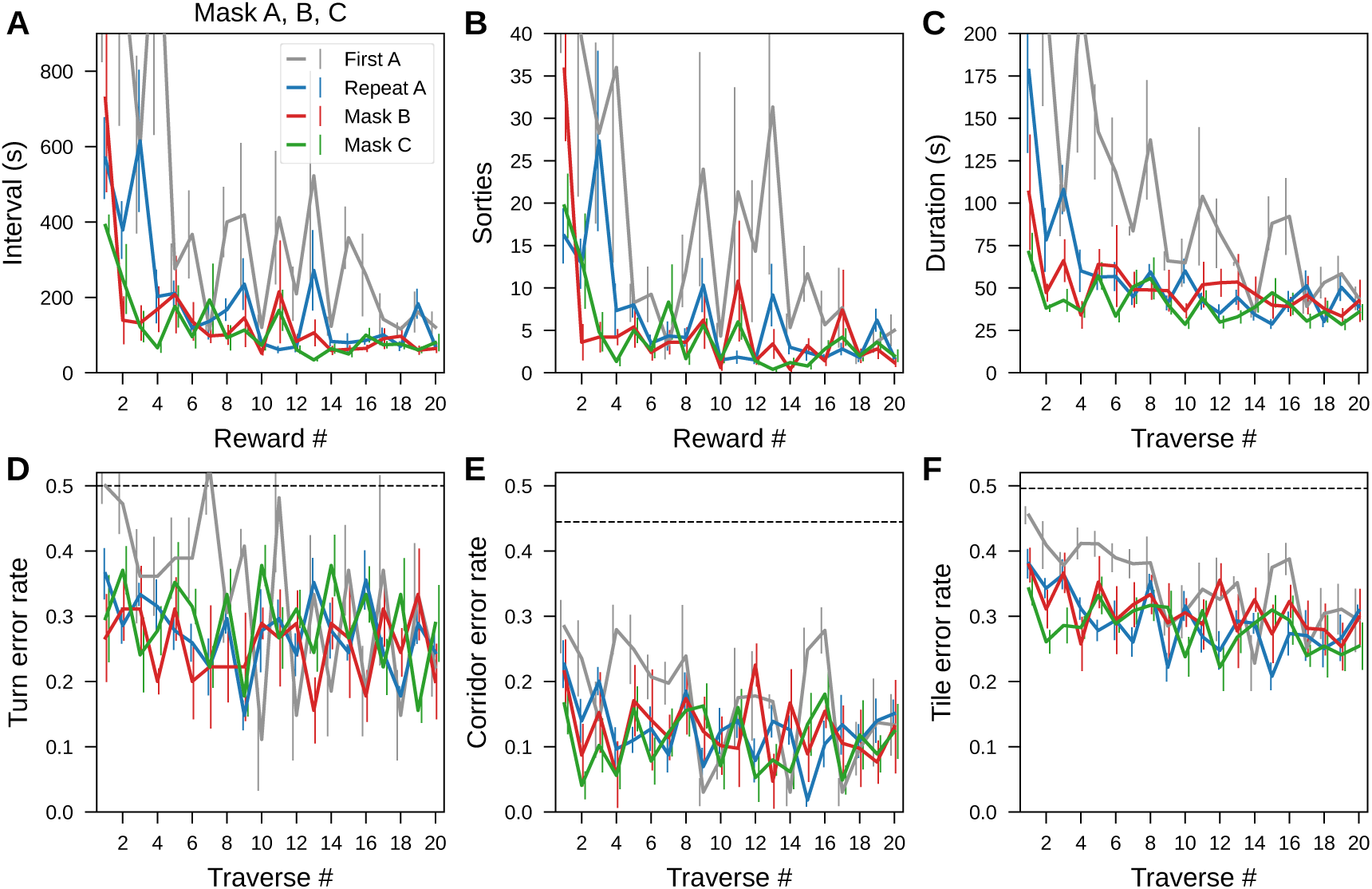
Acortical mice generalized to Masks B and C. **A.** Reward intervals on new masks (Masks B and C) and repeated Mask A, compared to the first time in the Mask A maze (gray). **B.** Number of sorties between rewards. Gray curve is identical to the orange curve in Fig. S8E. **C.** Traverse duration. Gray curve is identical to the orange curve in Fig. S8G. **D.** Turn error rate of traverses. Gray curve is identical to the orange curve in Fig. S8H. Dashed line, chance error rate (0.5). **E.** Corridor error rate of traverses. Dashed line, chance error rate (0.44). **F.** Tile error rate of traverses. Dashed line, chance error rate (0.50).

**Supplementary Figure S12:**
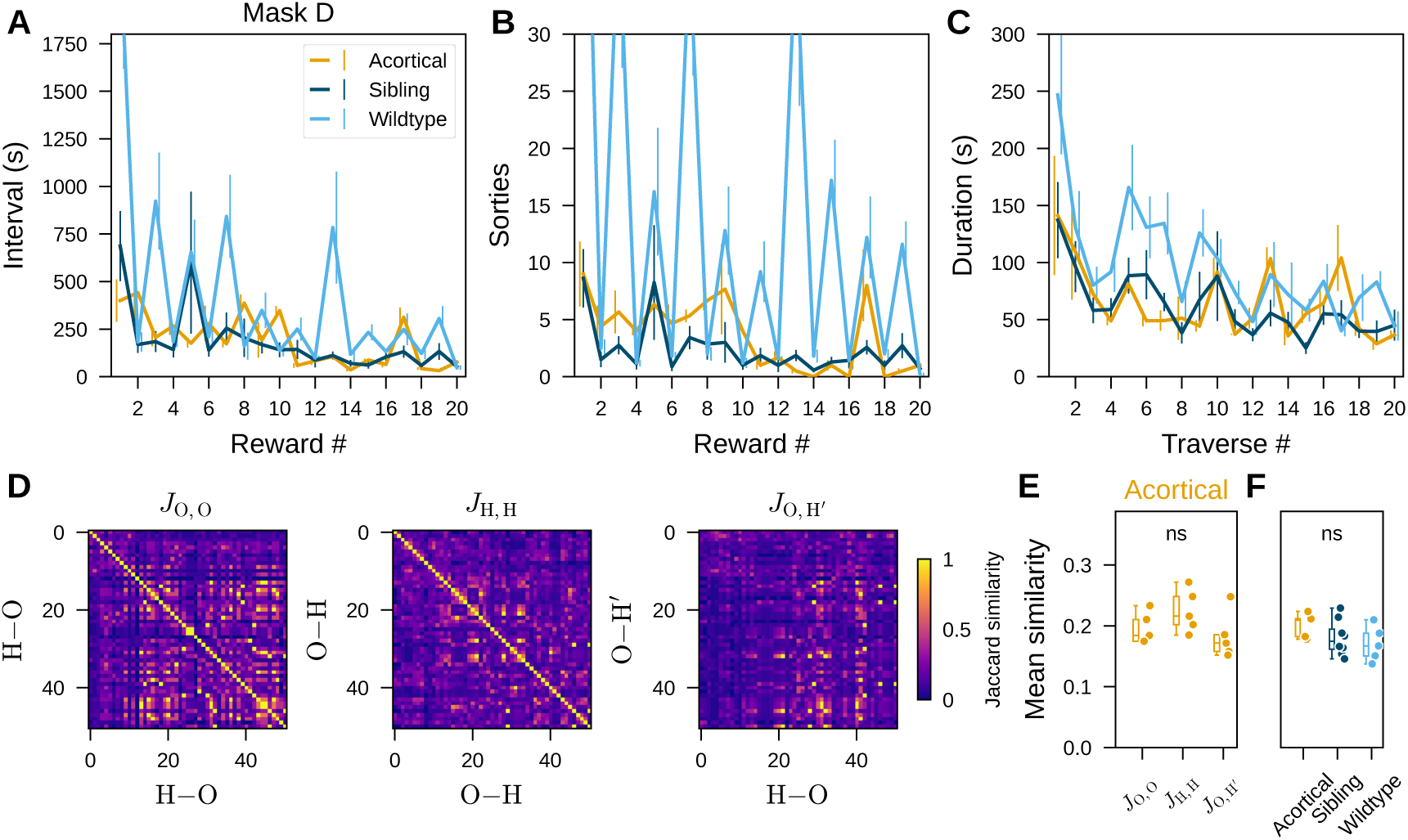
Acortical mice generalized to Mask D without repeating a fixed route. **A-C.** Mask D learning in acortical (5 mice, orange), sibling (8 mice, dark blue) and wildtype (6 mice, light blue); lines and error bars show mean ± SE across animals. **A.** Reward intervals. The light blue curve is the first 20 rewards of Fig. S4B. **B.** Number of sorties. The light blue curve is the first 20 rewards of Fig. S4C. **C.** Traverse duration. The light blue curve is the first 20 traverses of Fig. 3C. **D.** The three similarity matrices of one example acortical mouse, plotted as in Fig. S5A. **E.** Adjusted-Jaccard route similarity within and between traverse directions, acortical mice (5 mice; Friedman test not significant, *χ*^2^(2) = 5.20, *p* = 0.074). **F.** Mean route similarity per animal, compared across genotypes, restricted to animals scored in all three similarity groups (acortical 5, sibling 8, wildtype 5; Kruskal-Wallis test not significant, *H* = 2.44, *p* = 0.29).

**Supplementary Figure S13:**
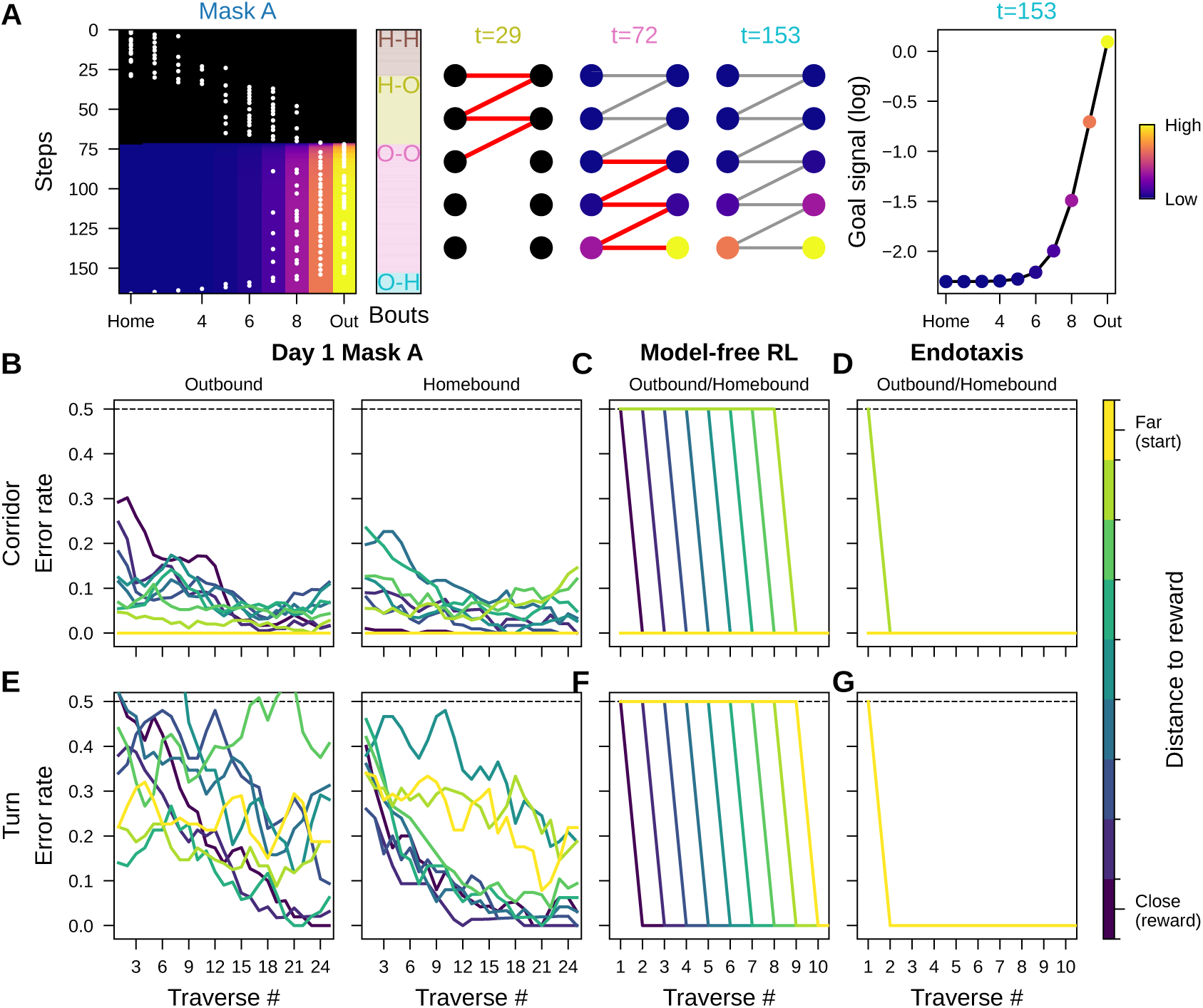
Mouse error propagation in the Mask A maze differs from model-free RL and Endotaxis predictions. **A.** Endotaxis model in the Mask A corridor graph, trained on the first two journeys of the mouse in Fig. 1E, as in Fig. 6. Left: propagation of the goal signal through corridor visits. Middle: goal signals and learned adjacency on the Mask A corridor graph, with newly added edges in red. Right: goal signal at the last timepoint (*t* = 153). **B-G.** Error rate propagation. Throughout, each line is one path position, colored by distance to reward from close (dark) to far (light); the dashed line marks chance (0.5). **B.** Corridor error rate per corridor across traverses, as the population mean over all 25 animals in Fig. 1G (smoothed with a 3-traverse moving-average window). **C.** Analytic prediction of the model-free RL agent for the same corridor error rate. **D.** Analytic prediction of the Endotaxis model for the same corridor error rate. **E-G.** As in **B-D**, for turn error rate per hole.

### 6.2 Supplementary videos

**Video V1. Manhattan Maze experiment setup.** Overview of the recording arena and task structure, followed by trimmed raw footage of the maze being reconfigured from Mask O to Mask A. The outbound and homebound traverses in the Mask O maze illustrate the mouse alternating between the two water ports (Fig. 1A and Fig. S1A-D). Shown at 2× speed.

**Video V2. First bouts of a wildtype mouse in the Mask A maze.** Continuous segment spanning the first 29 bouts of a wildtype mouse’s initial exposure to Mask A, including both traverses and sorties, with a scrolling inset of tile distance to the rewarded port (Fig. 1E). Shown at 8× speed.

**Video V3. Overnight memory in the Mask A maze.** Outbound and homebound traverses from Day 1 and Day 2.1 of the same mouse, each rendered individually with a growing inset of tile distance to reward colored to match the figure, illustrating overnight memory retention (Fig. 2C). Shown at 4× speed.

**Video V4. Late bouts in the Mask D maze.** Continuous segment of six consecutive traverses late in learning of the Mask D maze, including the interleaved sorties, with a bout trajectory inset (Fig. S5D). Shown at 8× speed.

**Video V5. Acortical mouse learning Mask A.** Continuous segment of an acortical mouse learning Mask A, including both traverses and sorties, with a scrolling inset of tile distance to the rewarded port (Fig. 4B). Shown at 8× speed.

**Video V6. Acortical and wildtype mouse late traverses in the Mask A maze.** The 19th and 20th Mask A traverses of an acortical mouse compared to the reference traverses of a wildtype mouse, each rendered individually with a growing inset of tile distance to reward colored to match the figure (Fig. 4D). The wildtype traverses are identical to the two in Video V3. Shown at 4× speed.

### 6.3 Maze construction and maintenance

The physical mazes in this study used a two-layered 11×11 structure (Fig. 1A and Fig. S1A). Each maze apparatus was made of clear acrylic (ePlastics, San Diego). The two layers were two identical, ∼18-inch square open-top trays of eleven 1.5-inch wide and 1.5-inch high corridors (wall thickness not included), with one access hole (1.5-inch diameter) on the side wall of the central corridor. We slotted the bottom plate (0.25 inches thick) of a tray every 1.5 inches, and then inserted ten walls (0.118 inches thick) into the slots to separate the tray into eleven corridors. The slots were laser cut to ensure that all eleven corridors were equally wide. The total width of the square tray was ∼18 inches, depending on the precise thickness of the materials. To form a maze, the two identical trays were stacked with bottom plates facing out, creating a closed space within the maze that could only be entered through the access holes. The top tray was rotated 90 degrees from the bottom tray, making the corridors in the two layers perpendicular. From the top view, the walls created an 11×11 grid pattern.

Between the two trays was a square mask with a series of holes. All masks were 0.125 inches thick, and the holes were of the same diameter (1.25 inches). Each hole was laser cut and centered at its corresponding grid coordinate. The masks were also made of clear acrylic, allowing imaging through the whole assembly via a top camera.

The mazes were sanitized regularly to remove odor and bacteria. The masks were wiped with disinfectants and deodorizers (Peroxigard, 1:16) before usage. After each day’s experiment, the maze trays were also cleaned thoroughly with deodorizers. All acrylic compartments contacted by the animals were submerged in room-temperature cleaning solution once per week. All items passed the annual NovaLum swab tests for sanitation verification.

### 6.4 Acortical mice learned intermediate masks and generalized

Because many acortical mice struggled with reward discovery in the more difficult masks, we tested whether they could learn an intermediate maze configuration. After learning Mask O, a subset of acortical mice was introduced to Mask E, a path graph with four corridors and three turns (Fig. S14A). These mice were initially unwilling to traverse through an additional hole and often became trapped in the vertical corridors. Therefore, they were guided through the holes until they connected movements across all corridors (Section 5.5.3). With this assisted training, 10 of 11 acortical mice learned Mask E (Table 2).

**Supplementary Figure S14:**
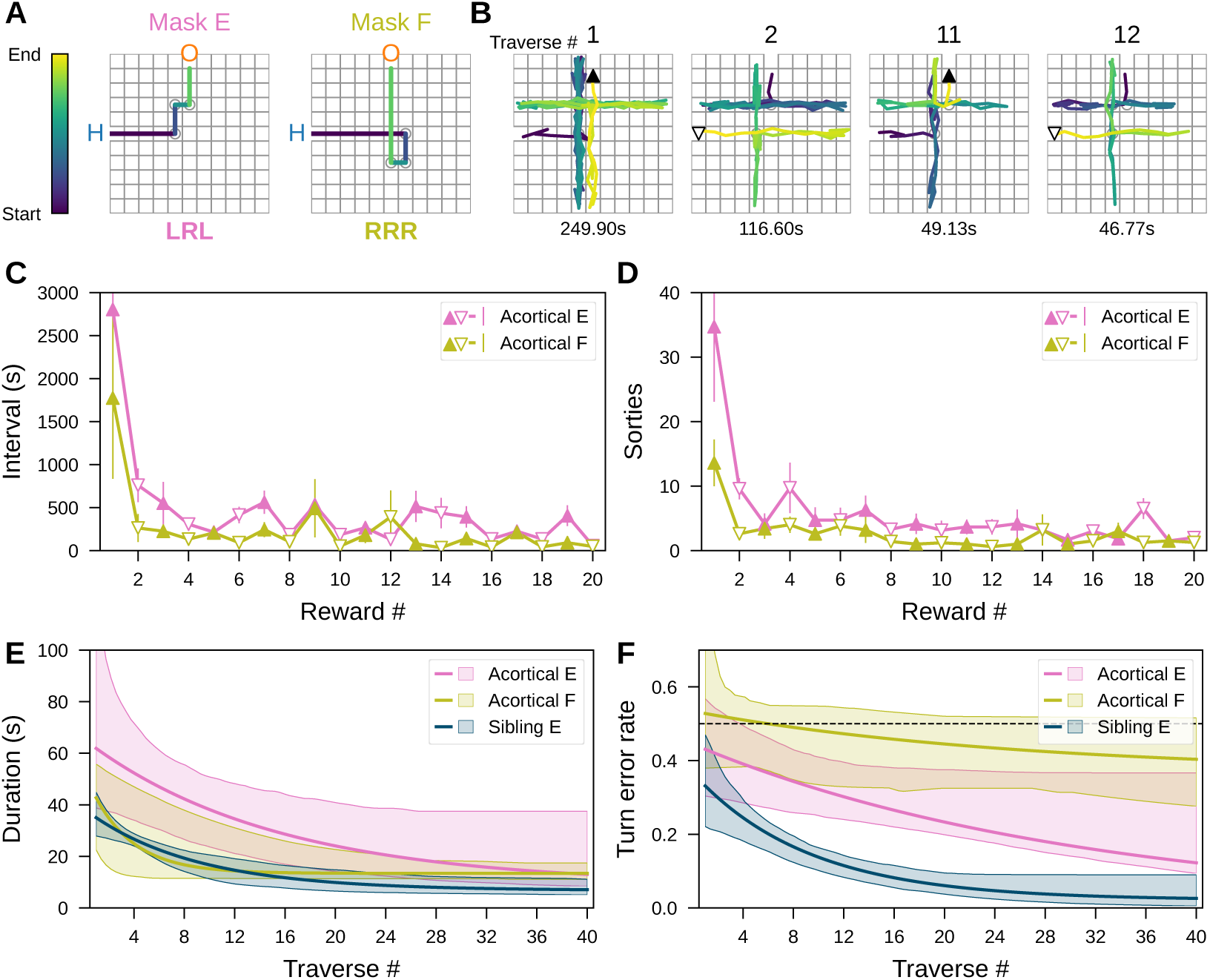
Acortical mice generalized from Mask E to Mask F. **A.** Top view of Mask E and Mask F. Of each mask, the shortest path is a sequence of three turns. **B.** Example traverses from an acortical mouse in the Mask E maze (text shows traverse duration). **C.** Reward intervals (session time) on Mask E (7 mice, pink) vs. Mask F (5 mice, olive, mean ± SE). **D.** Number of sorties between rewards (mean ± SE). **E.** Exponential fits of traverse duration for Mask E and Mask F, compared with sibling mice (6 mice, dark blue). **F.** Exponential fits of turn error rates of traverses. Dashed line, chance error rate.

Despite the simplicity of Mask E relative to Mask A, acortical mice still showed impaired initial performance. Their first traverses were prolonged by repetitive scanning (Fig. S14B,D). However, performance improved rapidly over the next 10 rewards: reward intervals shortened to ∼0.15 of the initial value (Fig. S14C). Consistent with the results of Mask A, acortical mice began with longer traverses than sibling mice but eventually reached similar asymptotic traverse durations (Fig. S14E), although their turn error rates remained higher than those of sibling mice (Fig. S14F).

We then tested whether acortical mice could transfer this experience to a related maze. Seven acortical mice were introduced to Mask F, a center-symmetric variant of Mask E (Fig. S14A). In contrast to their difficulty with Mask E, five mice learned Mask F without assistance. They obtained the first reward in ∼0.6 of the time required in the Mask E maze (Fig. S14C) and started with shorter traverse durations (Fig. S14E). Together, these results show that acortical mice can learn multiple maze configurations and that prior experience can facilitate learning of a related path graph.

### 6.5 Massive landmark changes did not reset turn learning

To test whether landmarks in the Manhattan Maze contribute to navigation, we performed a tray-swapping manipulation (Fig. S15A). If a mouse marks the maze with scent or urine, these deposits are likely to end up on the bottom tray or on the mask, so swapping the two trays and replacing the mask should severely disrupt such markers. In addition, any other markings, such as inadvertent construction glitches that identify a specific corridor, were perturbed as well.

Eight animals were first trained in the Mask O maze and then introduced to Mask A as in the main training protocol (Section 5.5.2). After each mouse obtained ∼20 rewards in the Mask A maze, we temporarily trapped it in the home cage, swapped the top and bottom trays of the two-layer maze, and inserted a clean copy of the Mask A plate. The home cage was then reconnected, allowing the mouse to resume exploration freely. The swap did perturb behavior transiently, but did not reset it: across the first ten post-swap traverses, turn errors were lower than in the first ten of the pre-swap session (Fig. S15B, methods from Section 5.6.5), most clearly for homebound turns, though they rose transiently over the first one or two post-swap traverses (Fig. S15C). One mouse did not enter the maze afterward, but 7 of 8 animals continued to obtain rewards (Fig. S15D). The swap thus did not produce a complete reset, and the animals did not have to relearn the turn sequences after the swap.

**Supplementary Figure S15:**
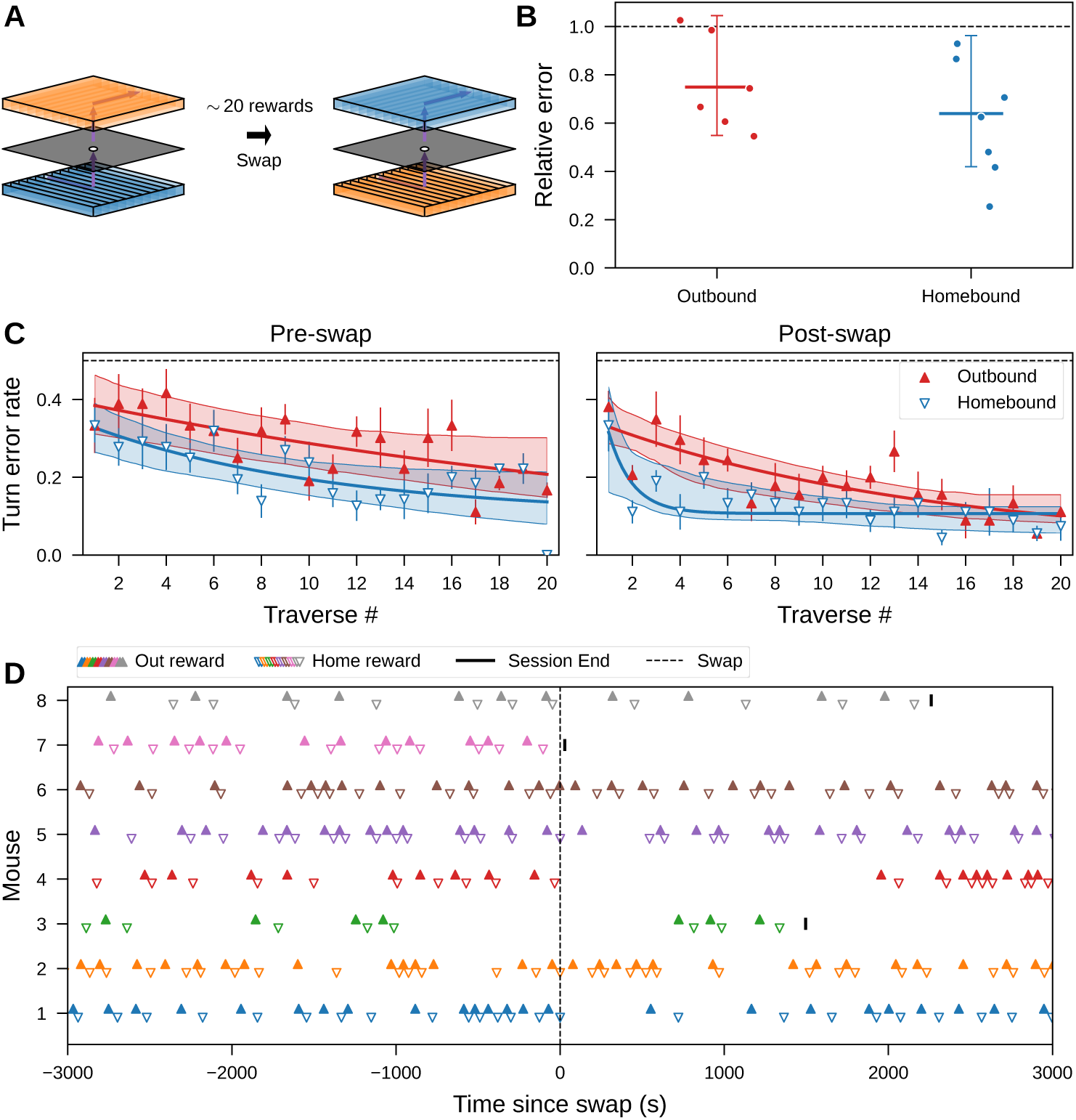
Mice quickly resumed learning after swapping the trays. **A.** The swapping procedure: after each mouse obtained ∼20 rewards in the Mask A maze, the top and bottom trays were exchanged and the mask was replaced with a clean, identical copy. **B.** Relative mean turn error rates of the first 10 traverses post-swap compared with the first 10 traverses in the pre-swap session, shown separately for outbound (red) and homebound (blue) directions (8 mice). The dashed line at 1 marks complete reset (values below 1 indicate savings). Error bars show 95% confidence intervals of the population-level mean (see text). Dots are individual animals. **C.** Outbound (red, filled triangles) and homebound (blue, open triangles) turn error rates before (left) and after (right) the swap. Points and error bars show population mean ± SE; solid lines are exponential fits of the population and shaded bands are their 95% bootstrap confidence intervals. Dashed line, chance error rate. **D.** Rasters of rewards of all mice in the experiment, aligned by the time (in maze) since the swap (dashed line at 0). Outbound (up triangles) and homebound (down triangles) rewards are shown per mouse; the pre-swap session runs in negative time and the post-swap session in positive time, and the solid line marks each session’s end.

**Supplementary Figure S16:**
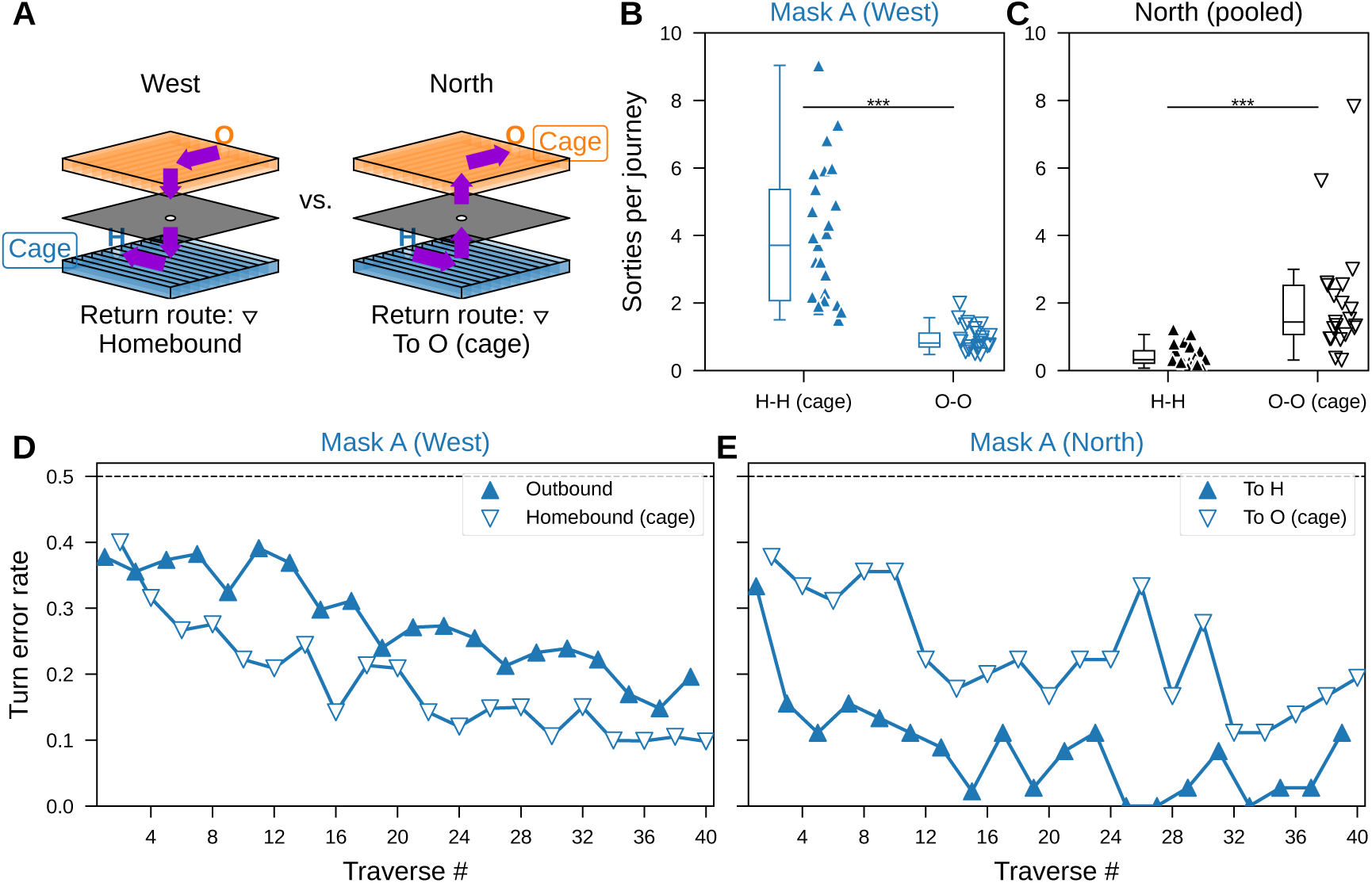
Maze navigation starting from O. **A.** The two start configurations. Left (“West”): the original layout in the main text, with the home cage on the bottom tray at the H port. Right (“North”): the cage relocated to the top tray at the O port, so that the return route is directed to O. **B.** Mean number of sorties per journey split by port (H-H and O-O) in the Day-1 Mask A session (“West” layout, Fig. 1; 25 sessions from 25 mice, same data as Fig. 1G). Asterisks show two-sided within-subject paired Wilcoxon signed-rank test (*W* = 0, *p* = 6 × 10*^−^*^8^). **C.** As in **B**, in the North layout (21 sessions pooled across 6 mice; two-sided Wilcoxon signed-rank *W* = 1, *p* = 1.9 × 10*^−^*^6^). **D.** Turn error rate in the Day-1 Mask A session (25 mice, mean ± SE). Each direction is connected as a separate line. Traverses toward H and cage are labeled “Homebound” (filled up-triangles) and traverses toward O are labeled “Outbound” (open down-triangles). Dashed line, chance error rate (0.5). **E.** Turn error rate in the Mask A maze under the North condition (5 mice, mean ± SE). Traverses toward H are labeled “To H” (filled up-triangles) and traverses returning to the cage at O are labeled “To O (cage)” (open down-triangles).

### 6.6 Outbound–homebound asymmetry was caused by route geometry

In the main experiments of Mask A, homebound traverses were less erroneous than the preceding outbound traverses (Fig. 1G). Two accounts could explain this asymmetry. First, the homebound direction might be learned faster because it is directed toward the home cage — a form of home-base behavior, in which rodents organize around the home base longer exploratory excursions away from home but shorter, more direct return trips [75]. Second, the two directions might simply be distinct physical routes whose geometry differs in difficulty, independent of cage location. In the Mask A maze, the outbound route requires climbing upward through 5 of 9 holes, whereas the homebound route descends. These accounts make opposite predictions when the cage is moved: a home-base advantage should follow the cage, so that the faster, more accurate traverses become those directed toward the new cage location, whereas a geometric advantage should remain bound to the same physical route regardless of where the cage sits.

To dissociate them, a new cohort was tested with the home cage moved from the H port to the O port while the mask orientation was held fixed. Mice therefore entered the maze via the top tray, from the “North” direction and, on each rewarded traverse, first traveled toward H — the physical route that had been the homebound route in the main experiments (Fig. S16A). If the asymmetry were motivational, performance should now favor the To-O traverses that return to the relocated cage; if geometric, it should still favor the H-directed route.

Home-base behavior itself was evident in the unrewarded sorties (Fig. S16B,C). In both layouts, mice launched more sorties from the cage-side port than from the opposite port, and this bias moved with the cage: from the H side under the West layout to the O side under North. On the other hand, the asymmetry tracked the physical route, not the cage. During initial training, mice hesitated to descend through the first hole but readily reversed and climbed once lowered to the bottom tray, after which the H-directed route was traversed fluently. In the Mask A maze, To-H traverses were less error-prone, consistent with the homebound traverses in the main text condition (Fig. S16D,E). Therefore, the outbound–homebound asymmetry in traverse performance in the main text is best explained by route geometry rather than by a home-base advantage.

